# Coordinating the morphogenesis-differentiation balance by tweaking the cytokinin-gibberellin equilibrium

**DOI:** 10.1101/2020.12.14.422551

**Authors:** Alon Israeli, Yogev Burko, Sharona Shleizer-Burko, Iris Daphne Zelnik, Noa Sela, Mohammad R. Hajirezaei, Alisdair R. Fernie, Takayuki Tohge, Naomi Ori, Maya Bar

## Abstract

Morphogenesis and differentiation are important stages in organ development and shape determination. However, how they are balanced and tuned during development is not fully understood. In the compound leaved tomato, an extended morphogenesis phase allows for the initiation of leaflets, resulting in the compound form. Maintaining a prolonged morphogenetic phase in early stages of compound-leaf development is dependent on delayed activity of several factors that promote differentiation, including CIN-TCP transcription factor (TF) LA, the MYB TF CLAU and the plant hormone Gibberellin (GA). Here, we investigated the genetic regulation of the morphogenesis-differentiation balance by studying the relationship between LA, CLAU and GA. Our genetic and molecular examination suggest that *LA* is expressed more broadly than *CLAU* and determines the spatio-temporal context of CLAU activity. We demonstrate that both LA and CLAU affect the Cytokinin/Gibberellin (CK/GA) balance. LA reduces the sensitivity of the leaf margin to CK, shown before to be also affected by CLAU. CLAU affects leaf active GA content and sensitivity, shown previously to be also influenced by LA. Therefore, LA and CLAU likely function in parallel pathways to promote leaf differentiation by converging on common downstream processes, including the CK/GA balance.

## Introduction

Morphogenesis, originating from the Greek words *morphe*/shape and *genesis*/formation, is a fascinating biological process that has attracted human eyes since ancient times (Theophrastus, 1916). Several model systems have been used to study morphogenesis, from the first examination of chicken embryos by Aristotle (Speybroeck *et al*., 2006; Munjal *et al*., 2015; Petit *et al*., 2017; Sutherland *et al*., 2020). Plants provide an excellent model system to investigate the shaping of an organism during the adult life cycle (Lintilhac, 2014; Pałubicki *et al*., 2019). Despite the ancient origin of morphogenesis studies in both the animal and plant kingdoms, our understanding of the molecular mechanisms governing morphogenesis, in particular the connection between gene regulatory networks, function, and shape formation - is still not complete.

Aristotle’s philosophy shaped our thinking of the term ‘form’ as fulfilling the full potential and destiny of oneself (Speybroeck *et al*., 2006). Leaves are vital photosynthetic, lateral organs produced by the plant throughout its life cycle. The development of plant leaves follows a common basic program, adjusted flexibly according to species, developmental stage and environment (Poethig, 1997; Bar & Ori, 2014; Chitwood & Sinha, 2016; Maugarny-Calès & Laufs, 2018). Morphogenesis and differentiation are important stages in leaf development, and the spatial and temporal balance between these processes influences leaf size and shape (Bar & Ori, 2014; Rodriguez *et al*., 2014; Du *et al*., 2018). In compound leaved plants such as tomato, the ratio between these two stages favors longer morphogenesis, allowing for initiation of leaflets, resulting in the compound form (Bar & Ori, 2015). The length of the morphogenetic window is thus a key determinant of final leaf shape. The flexibility of the morphogenetic window is regulated through a coordinated interplay between transcription factors and hormones (Raman *et al*., 2008; Shani *et al*., 2010; Busch *et al*., 2011; Yanai *et al*., 2011; Naz *et al*., 2013; Furumizu *et al*., 2015; Bar *et al*., 2016; Shwartz *et al*., 2016; Hajheidari *et al*., 2019). Tomato leaf development is therefore an attractive system to investigate the contribution of the morphogenesis-differentiation balance to organ shaping.

CIN-TCP transcription factors affect leaf shape by promoting differentiation, and maintenance of the morphogenetic window is dependent on low CIN-TCP activity during the early stages of leaf development (Nath *et al*., 2003; Palatnik *et al*., 2003; Ori *et al*., 2007; Efroni *et al*., 2008; Shleizer-Burko *et al*., 2011; Blein *et al*., 2013; Schommer *et al*., 2014; Ballester *et al*., 2015; Koyama *et al*., 2017; Challa *et al*., 2019). A subset of CIN-TCPs, including LANCEOLATE (LA) from tomato, is negatively regulated by the microRNA miR319. In the tomato semidominant gain-of-function mutant *La*, a mutation in the miR319 binding site leads to early ectopic LA expression, resulting in precocious differentiation and small, simplified leaves (Mathan & Jenkins, 1962; Dengler, 1984; Ori *et al*., 2007; Shleizer-Burko *et al*., 2011). Concurrently, premature expression of the miR319-insensitive *TCP4* in Arabidopsis plants causes early onset of maturation, resulting in a range of leaf patterning defects (Palatnik *et al*., 2003). Downregulation of *CIN-TCP* genes by overexpression of *miR319* results in a substantial delay in leaf maturation and prolonged indeterminate growth in the leaf margin (Koyama *et al*., 2007; Ori *et al*., 2007; Efroni *et al*., 2008; Shleizer-Burko *et al*., 2011; Challa *et al*., 2019). Differences in the timing of leaf growth and maturation among species and leaf positions are associated with altered *LA* expression dynamics (Shleizer-Burko *et al*., 2011). Thus, the *LA-* miR319 balance defines the morphogenetic window at the tomato leaf margin that is required for leaf elaboration. LA activity is mediated in part by positive regulation of the hormone gibberellin (GA) (Yanai *et al*., 2011).

Maintenance of the morphogenetic window is also restricted by activity of the MYB transcription factor *CLAUSA* (*CLAU*) (Avivi *et al*., 2000). CLAU has evolved a unique role in compound-leaf species to promote an exit from the morphogenetic phase of leaf development (Bar *et al*., 2016). *clau* mutants have highly compound, continuously morphogenetic leaves, in which meristematic tissues constantly generate leaflets on essentially mature leaves throughout the life of the plant (Bar *et al*., 2015, 2016). *clau* mutants can be extremely variable in phenotype, showing that tight regulation of the morphogenetic window is also required for shape robustness (Bar *et al*., 2015). CLAU regulates the morphogenetic window by attenuating cytokinin signaling and sensitivity (Bar *et al*., 2016).

While several transcription factors and hormones were shown to modulate the morphogenetic window during tomato leaf development, how their activities are coordinated is not clear. In this work, we investigate the relationship between the transcription factors LA and CLAU and the plant hormones GA and CK, in the regulation of the morphogenesis-differentiation balance. We show that LA and CLAU effect essentially similar outcomes in tomato leaf development via likely parallel pathways. They converge on modulation of the CK/GA balance.

## Results

### LA and CLA U operate in parallel pathways

To better understand the genetic regulation of the balance between morphogenesis and differentiation, we examined the interaction between the TFs CLAU and LA (**Figure 1** and (Ori *et al*., 2007; Bar *et al*., 2016). Increasing *CLAU* levels in a high *LA* expression background (*FIL>>CLAU La-2/+*) (**Figure 1D,M**) exacerbates the highly differentiated *La-2/+* phenotype (Figure **1B**). Interestingly, *FIL>>CLAU La-2/+* leaves were similar in size and complexity to *La-2* homozygous leaves (Compare **Figure 1D** and **Figure 6A**), suggesting an additive, dosedependent interaction. In agreement, decreasing *CLAU* levels in a low *LA* expression background such as *la-6* or *FIL>>miR319* results in a significant increase in leaf elaboration **(Figure 1G,K,N**). Decreasing *CLAU* levels in a high *LA* expression background (**Figure 1F,N**) partially rescues the highly differentiated *La-2/+* phenotype, while increasing *CLAU* levels in a low *LA* expression background (**Figure 1K,L**) results in a decrease in leaf elaboration when compared with the decreased *LA* expression genotypes. These genetic analyses demonstrate that the effect of *CLAU* and *LA* on leaf development is partially additive, indicating that they promote differentiation in parallel pathways.

### LA *activity defines the expression window of CLAU*

Our previous results demonstrated that *LA* has a wider expression window than *CLAU*, and is active earlier in development (Ori *et al*., 2007; Shleizer-Burko *et al*., 2011; Bar *et al*., 2016). Examination of the dynamics of *CLAU* expression in the first and fifth leaves of the plant, which represent a relatively limited and a relatively extended morphogenetic window, respectively (Shleizer-Burko *et al*., 2011), confirmed that *CLAU* is expressed mostly during the extended morphogenetic window (**Figure S1**). As the morphogenetic window is partially defined by LA (Ori *et al*., 2007; Shleizer-Burko *et al*., 2011), this raised the possibility that low LA activity enabled the recruitments of CLAU in the regulation of leaf differentiation. To explore this possibility and gain more insight into the molecular basis of the additive interaction of LA and CLAU in promoting differentiation, we examined how LA activity affects *CLAU* expression, by assaying the expression of *CLAU* and its promoter in successive stages of leaf development in genotypes with altered LA activity (**Figure 2**). Early maturation caused by increased *LA* expression in the gain-of-function mutant *La-2*, led to a decrease in *CLAU* expression (**Figure 2E-H,M**). Conversely, delayed maturation resulting from decreased *LA* expression in *FIL>>miR319* resulted in increased *CLAU* expression (**Figure 2 I-L,M**). Interestingly, expressing a miR319-resistant form of LA (*op:La-2*) from the *CLAU* expression domain mimicked the *La-2/+* phenotype (**Figure S2)**, with a slightly weaker effect when compared to expressing the same *La-2* version from its own expression domain (Burko *et al*., 2013). We conclude that *LA* activity defines the spatiotemporal expression domains for *CLAU*, which are limited in *La-2/+* and increased in *FIL>>miR319*. Therefore, these TFs act in partially distinct spatial and temporal domains to promote differentiation.

### Common TKN2-promoted pathways mediate extended morphogenesis

The class I KNOX homeobox transcription factor TKN2 is a key factor promoting morphogenesis that similarly to CLAU, was investigated in the context of compound leaf morphogenesis (Koltai & Bird, 2000; Hay *et al*., 2002; Grigg *et al*., 2005; Hay & Tsiantis, 2006, 2010; Kimura *et al*., 2008; Shani *et al*., 2009; Rast-somssich *et al*., 2015). Therefore, we set to examine the role of *TKN2* in mediating the extended morphogenesis in plants with reduced CLAU or LA activity, by combining them with expression of *TKN2-SRDX*, in which downstream targets of *TKN2* are inhibited (Shani *et al*., 2009). Expressing *TKN2-SRDX* from the *BLS* promoter lacks any observable phenotype in the WT background (Shani *et al*., 2009). Interestingly, *BLS>>TKN2-SRDX* suppresses the increased complexity of *CLAU* or *LA* deficient backgrounds (**Figure 3E, F, G**). In addition, reducing TKN2 targets from the *LA* expression domain (*LA>>TKN2-SRDX*) resulted in similar phenotypes to those of *La-2/+* mutants (**Figure S2B**), which stresses the important role of TKN2 downstream of LA. In contrast, expressing *TKN2* from either the *LA* or *CLAU* expression domains (*pLA>>TKN2* and *pCLAU>>TKN2*, respectively) produced highly compound leaves (**Figure S2C and D)**. This was more evident in the case of the *CLAU* promoter compared with the *LA* promoter. This is in accordance with *CLAU* being expressed later than *LA* during the morphogenetic window, and with the effect of stage-specific expression of *TKN2* (**Figure S2B, D** and (Ori *et al*., 2007; Shani *et al*., 2009; Shleizer-Burko *et al*., 2011; Burko *et al*., 2013; Bar *et al*., 2016)). Together, these results indicate that TKN2 mediates the increased complexity resulting from compromised CLAU and LA activities, and that *LA* and *CLAU* may have partially overlapping functions with *TKN2*. In agreement with previous findings (Avivi *et al*., 2000; Jasinski *et al*., 2007), the *TKN2* promoter is more strongly activated at the leaf margin of *CLAU* or *LA* deficient backgrounds than in the wild type, while remaining mostly restricted to meristems in WT. Its expression is further elevated in the *clau la-6* double mutant background (**Figure S3**). To investigate the functional interaction among these factors, we examined the effect of combining altered *CLAU* and *LA* expression with *TKN2* overexpression (**Figure 4**). Overexpressing *TKN2* (**Figure 4G**) in a *CLAU* (**Figure 4B**) or *LA* (**Figure 4D,E**) deficient background (**Figure 4 H,J,K,N**), leads to highly compound leaf forms. Overexpressing *TKN2* in a *CLAU* (**Figure 4C**) or *LA* (**Figure 4F**) overexpression background (**Figure 4 I,L,N**) leads to increased relative leaf elaboration and a rescue of the simplified leaf forms generated by overexpression of *CLAU* or *LA*. This rescue is substantial in the case of *CLAU*, and more moderate in the case of *La-2/+* (**Figure 4**). Terminal leaflets are exemplified in shading in Figure 3M. These results indicate that the phenotypes observed upon loss of function of *CLAU* or *LA* are not solely due to *TKN2* and that there are other morphogenesis-differentiation processes mediated by LA and CLAU.

### CLAU and LA interaction converges on the CK-GA balance

We previously demonstrated that *CLAU* functions through attenuation of CK signaling (Bar *et al*., 2016). We and others have also previously shown that LA functions in part through GA signaling (Yanai *et al*., 2011; Silva *et al*., 2019). Previous work has also demonstrated that the Arabidopsis *LA* homolog, *TCP4*, reduces leaf CK response through binding and promoting expression of the CK response inhibitor *ARR16* (Efroni *et al*., 2013). Since *CLAU* and *LA* exert similar developmental effects in different pathways and spatiotemporal windows, and since CK and GA are partially antagonistic in leaf development (Ori *et al*., 2007; Weiss & Ori, 2007; Fleishon *et al*., 2011; Bar *et al*., 2016), we examined the relationship between *CLAU* and GA, and *LA* and CK. In agreement with the antagonistic relationship between CK and GA in leaf morphogenesis, reducing CK content by overexpression of the CK inactivation gene CKX, or application of GA, led to simplification of leaf form (**Figure 5B,D**), and combining reduced cytokinin with increased GA further reduced leaf complexity (**Figure 5H**). Interestingly, the leaves of simultaneously reduced CK and increased GA levels resulted in phenotypes that were very similar to that of *La-2* and *CLAU* overexpression plants (**Figure 1)**. Conversely, inhibition of GA response via overexpression of a GA-resistant form of the GA response inhibitor DELLA/PROCERA (PRO) (PROΔ17) results in increased leaf complexity (**Figure 5E**). Interestingly, the simplified leaf phenotype caused by *CLAU* overexpression (**Figure 5C**) is rescued by co-expression of *PROΔ17* (**Figure 5G**), suggesting that GA may mediate the effect of CLAU on leaf differentiation.

To further understand the role of GA downstream of CLAU, we examined the effect *clausa* on GA content. Interestingly, GA4 and GA20 amounts were substantially reduced in 14-day-old *clausa* shoot apices, while the content of the more upstream GAs GA53 and GA19 increased (**Figure 5M**). This demonstrates that the GA pathway is altered in *clausa* mutants. In agreement with the accumulation of GA19 and decrease in GA20, and with previous findings (Jasinski *et al*., 2008), the expression of *SlGA20ox-1* was reduced in *clausa* mutants (**Figure 5N**). These results suggest that CLAU promotes differentiation by regulating GA biosynthesis, and that in *clau* mutants, reduced GA levels and/or response facilitate prolonged morphogenesis and compound leaf shape.

To examine whether CLAU also influences the leaf sensitivity to GA, we treated WT and *clausa* plants with increasing GA concentrations (Figure 5O). The *clausa* mutant displayed a strong and significant reduction in GA sensitivity at the leaf margin, remaining highly compound despite GA treatments at WT-responsive concentrations (0.01-1 uM GA), and responding only to a whopping 10 uM of GA (**Figure 5 I-L,O**). Therefore, CLAU exerts its role in regulating differentiation through regulation of both GA levels and response.

In kind, The *La-2/+* simple-leaf phenotype is exacerbated by overexpression of the CK inactivation gene CKX (*La-2/+ FIL>>CKX*) (**Figure 6**), as we reported previously for *CLAU* overexpression (Bar *et al*., 2016). In agreement, *La-2/+* is partially rescued by overexpression of CK biosynthesis gene *IPT (La-2/+ FIL>>IPT*) (**Figure 6E**). Reducing CK in a *LA* deficient background shortens the morphogenetic window, partially rescuing the super compound phenotype of *FIL>>miR319* (**Figure 6F**). In addition, similar to the arabidopsis TCP4, we found that LA reduced leaf sensitivity to CK (**Figure 6G-O and S5**). We found that LA, CLAU and the CK/GA balance also affect inflorescence complexity in a similar manner to their effect in the leaves (**Figure S6**). Thus we conclude that both CLAU and LA enhance differentiation by reducing the plant’s sensitivity to CK and by elevating GA levels and/or response. Together, LA and CLAU affect the GA/CK balance, in turn tuning the morphogenesis-differentiation balance.

### Global transcriptomic approach to identify common molecular pathways of morphogenesis and differentiation

To gain insights on leaf morphogenesis at the molecular level, we took a global transcriptomic approach. Our findings suggest that the key regulators: LA, CLAU and TKN2 act in partially parallel pathways that converge on the same downstream processes in the regulation of the balance between morphogenesis and differentiation. We thus compared several transcriptomic data sets from various genetic backgrounds with different activity of CLAU (WT vs *clausa*), LA (*La-2/+* gain-of-function, WT, *la-6* loss-of-function and *FIL>>miR319* that down regulates *LA* and three additional *CIN-TCPs: TCP3, TCP10* and *TCP24*) and TKN2 (*BLS>>TKN2* vs WT and *BLS>>TKN2-SRDX*) (**Figure 7**). Microarray data sets for the *LA* genotypes and *TKN2*, and RNAseq data for the *clausa* mutant, were analyzed for Fold change. KEGG (Kyoto Encyclopedia of Genes and Genomes) analysis was conducted to identify significantly differential pathways. Each genotype was compared to its WT background for the analysis. DEGs confirm dependencies between the LA genotypes, with between about a third to half of the genes significantly upregulated in *La-2/+* being significantly downregulated in *la-6* and upon *miR319* overexpression (**Figure 7**, **Supplementary Data 1**). Likewise, about a third of the genes significantly downregulated in *La-2/+* are significantly upregulated in *la-6* or upon *miR319* overexpression (**Figure 7**, **Supplementary Data 1 and 2**).

Interestingly, commonly up/downregulated genes are overrepresented between LA datasets and TKN2 datasets, with 2-3 times more DEGs than expected being commonly upregulated in *La-2/+* and downregulated upon *TKN2* overexpression or upregulated upon *TKN2-SRDX* overexpression (**Supplementary Data 1 and 2**). In agreement with our genetic and molecular analyses, genes upregulated upon low *LA* expression (*la-6, miR319* overexpression), or genes downregulated upon high *LA* expression (*La-2/+*), correlate best with those upregulated upon *TKN2* overexpression or downregulated upon *TKN2-SRDX* overexpression (**Figure 7D**). This demonstrates that, to a degree that is significantly higher than expected from random sampling, the extended morphogenesis upon absence of LA activity correlates with increased TKN2 activity, and with downregulation of processes which are affected by inhibition of TKN targets.

KEGG pathway analysis of DEGs in all samples revealed that, in agreement with the results of this study (**Figure 5, 6**) and with published data (Yanai *et al*., 2011; Bar *et al*., 2016), plant hormone signal transduction pathways are affected in *TKN2, CLAU* and *LA* genotypes (**Figure 7, Supplementary Data 1 and 2**). For example, GA signaling is altered in these genotypes, with *DELLA/PROCERA* upregulated in *clausa* and the GA-receptor *GID1* upregulated in *miR319* and *TKN2* and downregulated in *La-2/+* and *TKN2-SRDX*, providing molecular context for the altered sensitivity of these genotypes to GA. Interestingly, jasmonate pathways are upregulated in all the “moprphogenetic” genotypes, most strongly in *clausa*, and ethylene signaling is uniquely upregulated in *clausa*, in additional to upregulation of plant pathogen interaction (p=0.00058) and MAPK signaling (p=0.0096) pathways (**Supplementary Data 1 and 2**). We have previously demonstrated that *clausa* is immuno-active and pathogen resistant (Gupta *et al*., 2020).

Analysis of increased morphogenesis genotypes (*la-6, miR319* overexpression, *clausa* and *TKN2* overexpression) revealed a significant increase in metabolic processes, carbon fixation, biosynthesis of amino acids and glycolysis - perhaps required for the increase in morphogenesis (**Supplementary Data 1 and 2**). Furthermore, it emerges that *LA* and *TKN2* co-regulate protein processing and protein modification, with pathways of ER protein processing being upregulated in *la-6* and *TKN2* and downregulated in *La-2/+*, and Glycosylphosphatidylinositol (GPI)-anchor biosynthesis being upregulated in *la-6* and *miR319* overexpression, and down regulated in *La-2/+* and *TKN2-SRDX* (**Supplementary Data 1 and 2**).

We compared our transcriptomic data to published data (Ichihashi *et al*., 2014) that includes three solanum species at four developmental stages. In the public data we focused on genes that showed successive downregulation throughout the developmental stages in the M82 background, which we termed ‘morphogenesis genes’, and on genes that were successively upregulated throughout the developmental stages in the M82 background, to whom we referred as ‘differentiation genes’. When comparing these sets of ‘morphogenesis’ and ‘differentiation’ genes with the DEGs in our genotypes we found that, in nearly all cases, morphogenesis genes were significantly enriched in morphogenetic genotypes *clausa*, *la-6*, *miR319* and *TKN2* over expressions, while differentiation genes were significantly depleted in these genotypes (**Figure 7, Supplementary Data 3 and 4**). In agreement, the differentiation genes were significantly enriched in *La-2/+* and depleted in the morphogenetic genotypes (**Supplementary Data 3 and 4**). Interestingly, the morphogenetic genes upregulated in *clausa* and *miR319* overexpression showed no overlap, while the differentiation genes depleted in *clausa* and *miR319* overexpression showed only 10% overlap, supporting the hypothesis that emerges from our results, that *CLAU* and *LA* may regulate different genetic mechanisms in the leaf developmental program (**Figure 7**).

## Discussion

Investigating the underlying molecular mechanism of shape formation is crucial for our understanding of organ function. In this work, we examined how the key regulators LA and CLAU interact in promoting and tuning differentiation during leaf development. Analysis of the interaction between *CLAU* and *LA* indicates that they operate in parallel pathways and suggests that LA might determine the window of *CLAU* activity (**Figures 1, S1**) (Shleizer-Burko *et al*., 2011). This is in agreement with the unique role of CLAU in compound leaf species (Bar *et al*., 2016). It will be interesting to identify additional compound-leaf specific regulators that are recruited in the context of extended morphogenesis. Similarly, the class I KNOX homeobox transcription factor TKN2 was also investigated in the context of extended morphogenesis of compound leaves (Koltai & Bird, 2000; Hay *et al*., 2002; Grigg *et al*., 2005; Hay & Tsiantis, 2006, 2010; Kimura *et al*., 2008; Shani *et al*., 2009; Rast-somssich *et al*., 2015), and was shown here to partially mediate the effect of both LA and CLAU in the regulation of the morphogenesis-differentiation balance. Here, we show that the CK-GA balance is a common process that mediates both LA and CLAU activity. Leaf development is known to depend on the balance between CK, which promotes morphogenesis, and GA, which promotes differentiation (Hay *et al*., 2002; Jasinski *et al*., 2005; Yanai *et al*., 2005; Shani *et al*., 2010; Fleishon *et al*., 2011; Scofield *et al*., 2013). The genetic interaction shown here between LA and CK (**Figure 6**), and previous reports showing an effect of TCP on the sensitivity to CK (Efroni *et al*., 2013), suggest that LA acts in part by reducing CK sensitivity. Previously, LA differentiation-promoting activity was shown to also depend on GA response (Maltnan & Jenkins, 1962; Yanai *et al*., 2011; Silva *et al*., 2019). In turn, CLAU promotes differentiation by elevating GA levels, and, in its absence, the plant becomes less sensitive to GA treatment at the leaf margin (**Figure 5**). CLAU was previously shown to act by reducing CK sensitivity (Bar *et al*., 2016). Therefore, CLAU and LA appear to converge on the CK-GA balance: both promote differentiation by increasing the plants’ response to GA and reducing its response to CK. The length of the morphogenetic window within leaf differentiation can thus be viewed as an almost binary “lever” of sorts: pulling the lever towards CK will lengthen the window, while pulling it towards GA will shorten the window. It seems that the differentiation-morphogenesis and CK-GA balances are regulated and interpreted in a dose dependent manner (**Figure 8**). The mutation in the miR319 recognition site in *La-2* is dominant, with the homozygote being more severely affected than the heterozygote (Maltnan & Jenkins, 1962; Ori *et al*., 2007). Our results demonstrate a “gradient” of transcription factor activity and hormone levels that is translated to leaflet number. Overexpression of both CLAU and LA, or either one of these transcription factors overexpressed with CKX (**Figure 1, 6**; Bar et al 2016), or the homozygous version of the dominant *La-2* mutant, all exhibit simple leaves without any leaflets, indicating that the capacity for morphogenesis is embodied in the activity of LA, CLAU, CK and GA, acting in concert. It may suggest that additional regulators that were co-opted into the developmental program of compound leaves are regulating this balance. For example, KNOXI proteins such as *TKN2* regulate the CK-GA balance, by negatively regulating the expression of the GA biosynthesis gene *GA20oxidase* (*GA20ox*) and positively regulating the GA deactivation gene *GA2oxidase* (*GA2ox*) (Sakamoto *et al*., 2001; Hay *et al*., 2002; Jasinski *et al*., 2005; Bolduc & Hake, 2009). KNOXI proteins also activate CK biosynthesis genes and promote CK accumulation (Sakamoto *et al*., 2001; Jasinski *et al*., 2005; Yanai *et al*., 2005). Here we show that *GA20ox-1* is positively regulated by CLAU (**Figure 5**). It is therefore possible that the regulation of the CK-GA balance by CLAU and LA may be mediated in part through pathways common with *TKN2*. The GA-CK balance also plays a key role in meristem maintenance, which highlights the similarities between the shoot apical meristem and the transient meristematic phase that the leaf primordia in preserving and enabling organogenesis (Floyd & Bowman, 2010).

Figure 8 details a model depicting the roles of CLAU, LA and TKN2 in the CK/GA balance during leaf development. Both LA and CLAU may promote differentiation via inhibition of TKN2, though they also appear to have TKN2 independent activity. The activity of different transcription factors may affect the location of the lever between CK and GA and can do so within different spatial-temporal domains of the developmental program.

Overall, The genetic, molecular, and transcriptomic analyses we present here, provide insights into the molecular basis of differentiation and morphogenesis processes in plants, that will be interesting to examine in the future in more species and developmental processes.

## Materials and Methods

### Plant Material

Tomato seeds (*Solanum lycopersicum* cv M82 or as indicated) were sown in a commercial nursery and grown in the field or in a glasshouse under natural daylight with 25:18°C (day: night) temperatures and a maximum light intensity of 450 μmol m^−2^ s^−1^. For developmental and expression analyses, plants were grown in a controlled growth chamber, 300 μmol m^−2^ s^−1^ 18 h/6-h light/dark regime.

Genotypes used in the present study were previously described: *clausa* (Menda *et al*., 2004; Bar *et al*., 2015, 2016). *pFIL>>CLAU* (Bar *et al*., 2016). *La-2/+* and *pFIL>>miR319* (Ori *et al*., 2007; Shleizer-Burko *et al*., 2011). *pFIL>>IPT* and *pFIL>>CKX* (Shani *et al*., 2010). *pBLS>>TKN2 and pBLS>> TKN2-SRDX* (Shani *et al*., 2009)*. pFIL>>PROΔ17* (Nir *et al*., 2017). *pTKN2::nYFP* was generated by amplifying ~5500 bp of genomic DNA upstream to the tomato *TKN2* atg using the primers detailed in Supplemental Table 1, fusing them to YFP with a nuclear localization signal, and transforming tomato plants – essentially as previously described for *pCLAU::nYFP* (Bar *et al*., 2016). Additional genotypes were generated by crossing these genotypes, where indicated. *pTKN2::nYFP, pCLAU::nYFP* (Bar *et al*., 2016), and *pTCSv2:3XVENUS* (Zürcher *et al*., 2013; Bar *et al*., 2016; Steiner *et al*., 2016), were backcrossed into the relevant backgrounds.

### Tissue Collection and RNA Analysis

Tissue collection, RNA preparation, and qRT-PCR analysis were performed as previously described (Shleizer-Burko *et al*., 2011). Expression of all assayed genes was normalized relative to tomato EXPRESSED (EXP). Primer sequences used in qRT-PCR analyses are detailed in Supplemental Table 1.

### Imaging

Leaves were photographed using a Nikon D5200 camera. For analysis of *pTKN2::nYFP, pCLAU::nYFP*, and *pTCSv2:3XVENUS* expression, dissected whole-leaf primordia were placed into drops of water on glass microscope slides and covered with cover slips. The pattern of YFP or VENUS expression was observed using a confocal laser scanning microscope (CLSMmodel SP8; Leica), with a solid-state laser set at 514 nm for excitation/ 530 nm for emission. Chlorophyll expression was detected at 488 nm excitation/ 700 nm for emission.

### GA Content Analysis

Giberellins were isolated and purified according to the method described by (Šimura *et al*., 2018).

### Anthocyanin Measurement

For anthocyanin measurement, plants were sprayed with the indicated CK concentrations three times a week for 3 weeks prior to analysis, starting upon emergence of the first leaf. Anthocyanins were extracted from the terminal leaflet of the third leaf by incubation overnight in methanol supplemented with a final concentration of 1% HCl. OD was measured in a plate spectrophotometer and anthocyanin content was calculated according to the following formula: (OD530(0.25*OD660)), normalized to the starting tissue weight. Three technical replicates of 5 8 biological repeats were performed for each sample.

### Electrophoresis Mobility Shift Assay (EMSA)

DNA probes were generated by end labeling of a 60-base single-stranded oligonucleotide using the DNA 3’ End Biotinylation Kit (Pierce 89818) and hybridization to complementary synthetic oligonucleotides (Supplemental Table 1) spanning binding sites for LA (GGNCC) which were identified using Sequencer 4.9, and generated with mutations disrupting the binding sites in the case of the mutant probe. Probes were generated by hybridizing the two complementary oligos by boil/cool. EMSAs were performed using the Light-Shift chemiluminescent EMSA kit (Pierce 20148). Briefly, 10 μL of purified recombinant MBP-LA fusion protein was incubated at room temperature in 1× binding buffer, 50 ng/μL poly(dI/dC), 2.5% glycerol, 0.05% Nonidet P-40, 50 fmol biotin-labeled probe, and 3.75 μg BSA for 30 to 40 min. The samples were resolved on 6% nondenaturing polyacrylamide gels, electrotransferred onto 0.45 μm Biodyne B nylon membrane (Pierce 7701), and cross-linked to the membrane. The migration of the biotin-labeled probe was detected on x-ray film (5-h exposure) using streptavidin–horseradish peroxidase conjugates and chemiluminescent substrate according to the manufacturer’s protocol.

## Supporting information

description of supplemetal datasets

Supp dataset 1- Lists of upregulated DEGs in all genotypes

Supp dataset 2- Differential KEGG pathways in all genotypes

Supp dataset 3- morphogenetic and diff genes from Ichihashi 2014

Supp dataset 5 - KNOX putative targets integrated with Ichihashi 2014

Supp dataset4- morphogenesis and differentiation

---

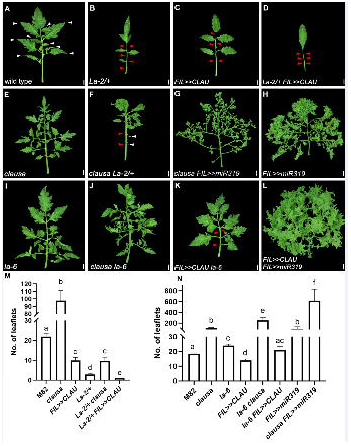

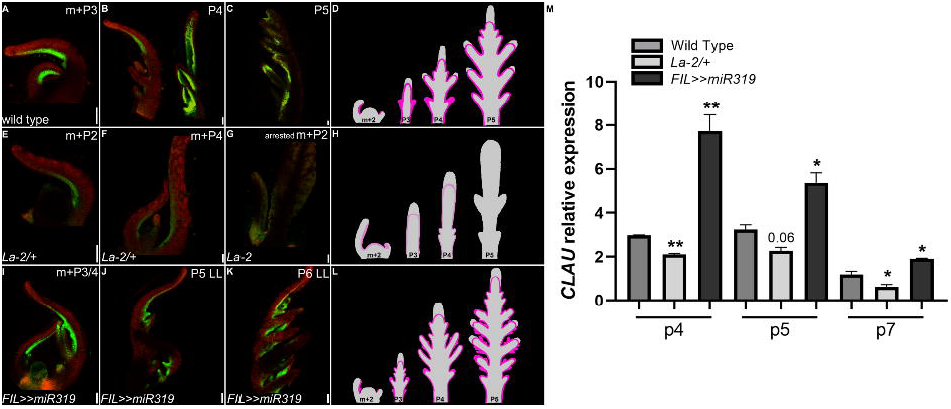

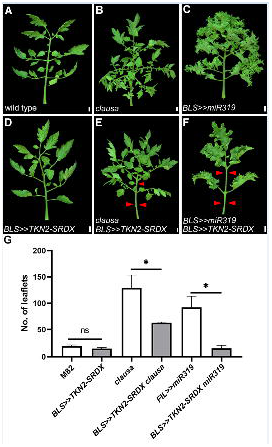

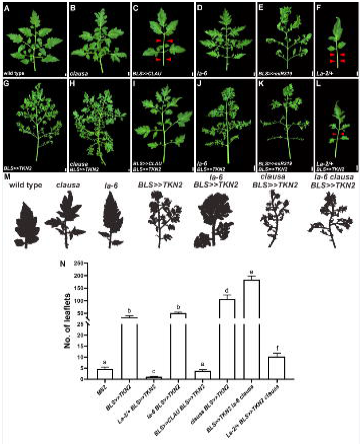

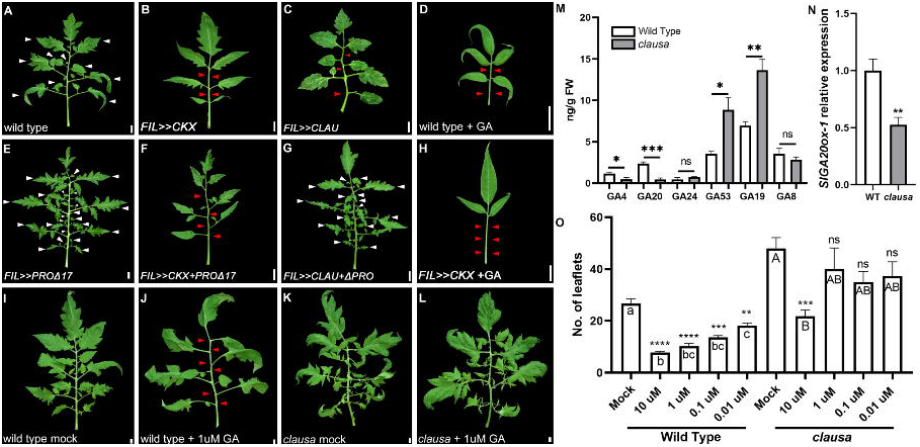

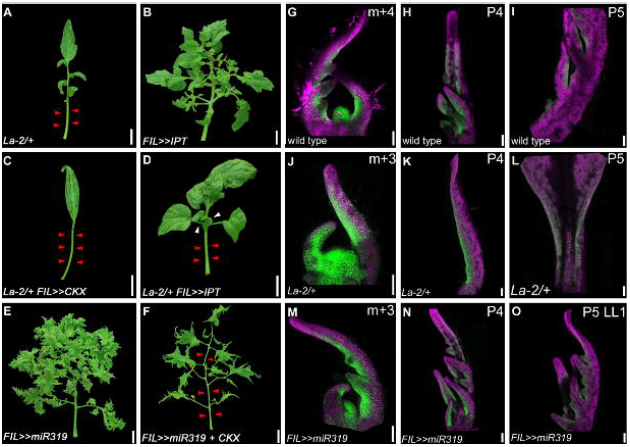

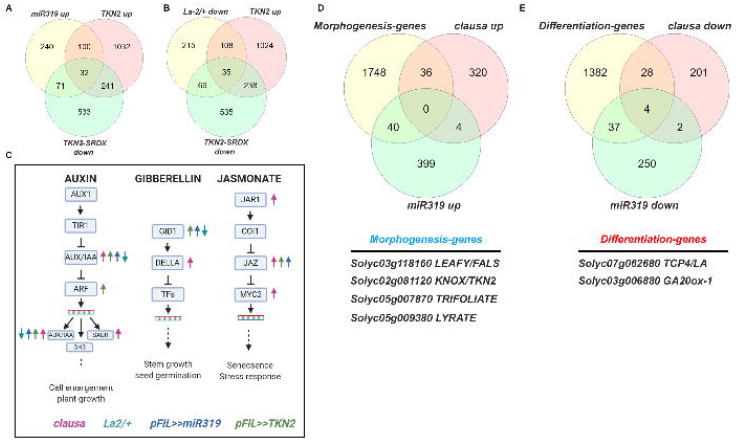

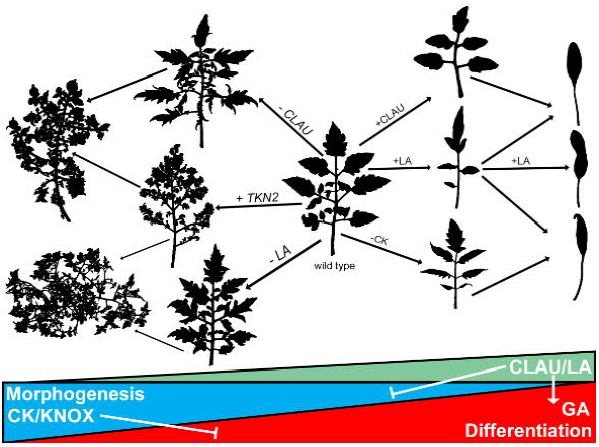

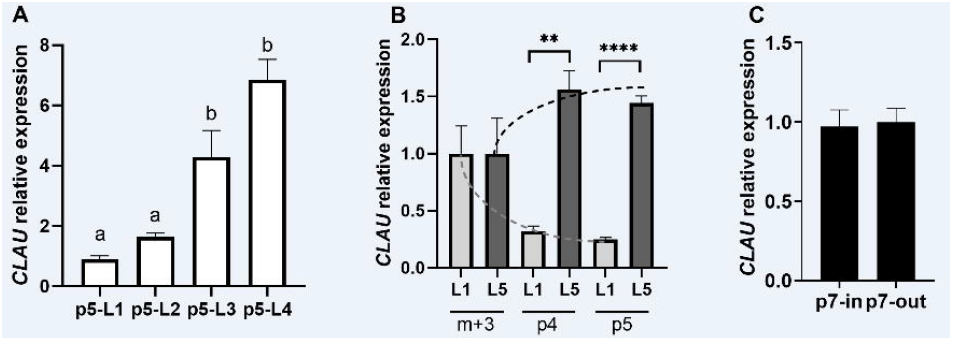

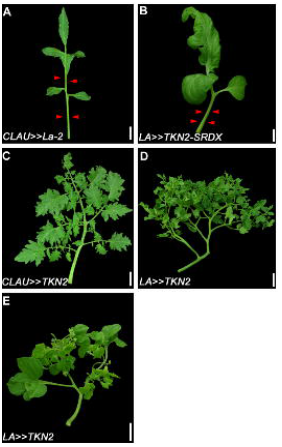

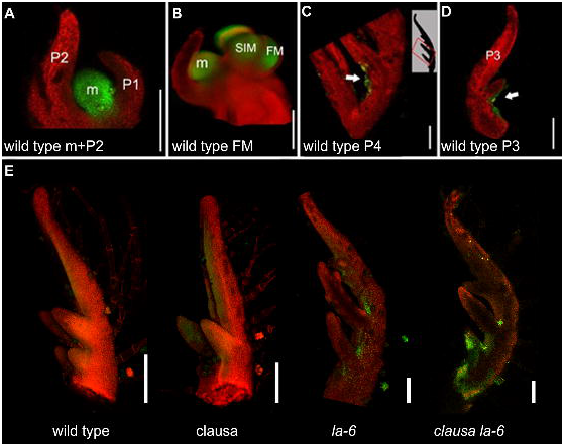

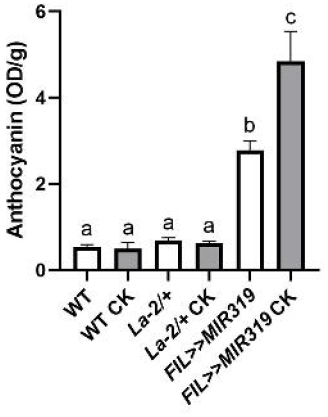

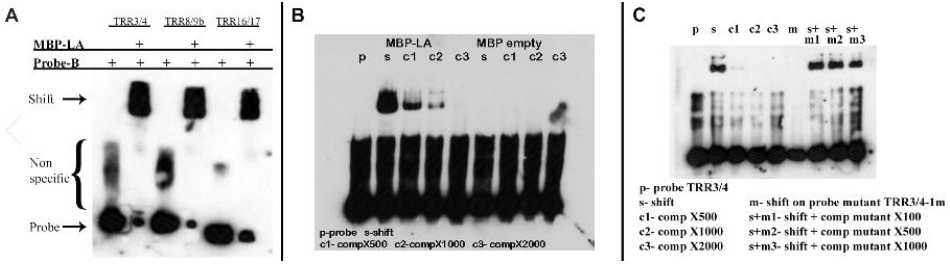

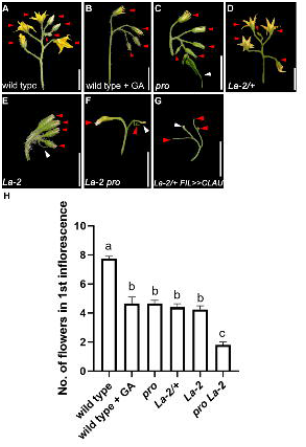

## References

Avivi Y, Lev-yadun S, Morozova N, Libs L, Williams L, Zhao J, Varghese G, Grafi G. 2000a. Clausa, a Tomato Mutant with a Wide Range of Phenotypic Perturbations, Displays a Cell Type-Dependent Expression of the Homeobox Gene LeT6 / TKn2 1. Plant physiology 124: 541–551.

Avivi Y, Lev-Yadun S, Morozova N, Libs L, Williams L, Zhao J, Varghese G, Grafi G. 2000b. Clausa, a tomato mutant with a wide range of phenotypic perturbations, displays a cell type-dependent expression of the homeobox gene LeT6/TKn21. Plant Physiology 124: 541–551.

Ballester P, Navarrete-Gomez M, Carbonero P, Onate-Sanchez L, Ferrandiz C. 2015. Leaf expansion in Arabidopsis is controlled by a TCP-NGA regulatory module likely conserved in distantly related species. Physiologia Plantarum 155: 21–32.

Bar M, Ben-Herzel O, Kohay H, Shtein I, Ori N. 2015. CLAUSA restricts tomato leaf morphogenesis and GOBLET expression. The Plant journal□: for cell and molecular biology 83: 888–902.

Bar M, Israeli A, Levy M, Ben Gera H, Jiménez-Gómez J, Kouril S, Tarkowski P, Ori N. 2016. CLAUSA is a MYB Transcription Factor that Promotes Leaf Differentiation by Attenuating Cytokinin Signaling. The Plant Cell 28: 1602–1615.

Bar M, Ori N. 2014. Leaf development and morphogenesis. Development (Cambridge, England) 141: 4219–30.

Bar M, Ori N. 2015. Compound leaf development in model plant species. Current opinion in plant biology 23: 61–9.

Blein T, Pautot V, Laufs P. 2013. Combinations of Mutations Sufficient to Alter Arabidopsis Leaf Dissection. Plants 2: 230–247.

Bolduc N, Hake S. 2009. The maize transcription factor KNOTTED1 directly regulates the gibberellin catabolism gene ga2ox1. The Plant cell 21: 1647–58.

Burko Y, Shleizer-Burko S, Yanai O, Shwartz I, Zelnik ID, Jacob-Hirsch J, Kela I, Eshed-Williams L, Ori N. 2013. A role for APETALA1/fruitfull transcription factors in tomato leaf development.

Busch BL, Schmitz G, Rossmann S, Piron F, Ding J, Bendahmane A, Theres K. 2011. Shoot branching and leaf dissection in tomato are regulated by homologous gene modules. The Plant cell 23: 3595–609.

Challa KR, Rath M, Nath U. 2019. The CIN-TCP transcription factors promote commitment to differentiation in Arabidopsis leaf pavement cells via both auxin-dependent and independent pathways. PLoS Genetics 15: 1–30.

Chitwood DH, Sinha NR. 2016. Evolutionary and Environmental Forces Sculpting Leaf Development. Current Biology 26: R297–R306.

Dengler NG. 1984. Comparison of Leaf Development in Normal (+/+), Entire (e/e), and Lanceolate (La/+) Plants of Tomato, Lycopersicon esculentum ‘Ailsa Craig’. Botanical Gazette 145: 66–77.

Du F, Guan C, Jiao Y. 2018. Molecular Mechanisms of Leaf Morphogenesis. Molecular plant 11: 1117–1134.

Efroni I, Blum E, Goldshmidt A, Eshed Y. 2008. A Protracted and Dynamic Maturation Schedule Underlies Arabidopsis Leaf Development. the Plant Cell Online 20: 2293–2306.

Efroni I, Han S-K, Kim HJ, Wu M-F, Steiner E, Birnbaum KD, Hong JC, Eshed Y, Wagner D. 2013. Regulation of Leaf Maturation by Chromatin-Mediated Modulation of Cytokinin Responses. Developmental Cell 24: 438–445.

Fleishon S, Shani E, Ori N, Weiss D. 2011. Negative reciprocal interactions between gibberellin and cytokinin in tomato. The New phytologist 190: 609–17.

Floyd SK, Bowman JL. 2010. Gene expression patterns in seed plant shoot meristems and leaves: Homoplasy or homology? Journal of Plant Research 123: 43–55.

Furumizu C, Alvarez JP, Sakakibara K, Bowman JL. 2015. Antagonistic Roles for KNOX1 and KNOX2 Genes in Patterning the Land Plant Body Plan Following an Ancient Gene Duplication. PLoS Genetics 11(2): e1004980.

Grigg SP, Canales C, Hay A, Tsiantis M. 2005. SERRATE coordinates shoot meristem function and leaf axial patterning in Arabidopsis. Nature 437: 1022–1026.

Gupta R, Pizarro L, Leibman-Markus M, Marash I, Bar M. 2020. Cytokinin response induces immunity and fungal pathogen resistance, and modulates trafficking of the PRR LeEIX2 in tomato. Molecular Plant Pathology 21: 1287–1306.

Hajheidari M, Wang Y, Bhatia N, Huijser P, Gan X, Tsiantis M. 2019. Autoregulation of RCO by Low-Affinity Binding Modulates Cytokinin Action and Shapes Leaf Article. Current Biology 29: 1–10.

Hay A, Kaur H, Phillips A, Hedden P, Hake S, Tsiantis M. 2002. The gibberellin pathway mediates KNOTTED1-type homeobox function in plants with different body plans. Current Biology 12: 1557–1565.

Hay A, Tsiantis M. 2006. The genetic basis for differences in leaf form between Arabidopsis thaliana and its wild relative Cardamine hirsuta. Nature Genetics 38: 942–947.

Hay A, Tsiantis M. 2010. KNOX genes: versatile regulators of plant development and diversity. Development (Cambridge, England) 137: 3153–3165.

Ichihashi Y, Aguilar-Martínez JA, Farhi M, Chitwood DH, Kumar R, Millon LV, Peng J, Maloof JN, Sinha NR. 2014. Evolutionary developmental transcriptomics reveals a gene network module regulating interspecific diversity in plant leaf shape. Proceedings of the National Academy of Sciences of the United States of America 111: E2616–E2621.

Jasinski S, Kaur H, Tattersall A. 2007. Negative regulation of KNOX expression in tomato leaves. Planta 226: 1255–1263.

Jasinski S, Piazza P, Craft J, Hay A, Woolley L, Rieu I, Phillips A, Hedden P, Tsiantis M, Regu- CAR. 2005. KNOX Action in Arabidopsis Is Mediated by Coordinate Regulation of Cytokinin and Gibberellin Activities. 15: 1560–1565.

Jasinski S, Tattersall A, Piazza P, Hay A, Martinez-Garcia JF, Schmitz G, Theres K, McCormick S, Tsiantis M. 2008. PROCERA encodes a DELLA protein that mediates control of dissected leaf form in tomato. The Plant journal□: for cell and molecular biology 56: 603–12.

Kimura S, Koenig D, Kang J, Yoong FY, Sinha N. 2008. Natural Variation in Leaf Morphology Results from Mutation of a Novel KNOX Gene. Current Biology 18: 672–677.

Koltai H, Bird DMK. 2000. Epistatic repression of PHANTASTICA and class 1 KNOTTED genes is uncoupled in tomato. Plant Journal 22: 455–459.

Koyama T, Furutani M, Tasaka M, Ohme-Takagi M. 2007. TCP transcription factors control the morphology of shoot lateral organs via negative regulation of the expression of boundary-specific genes in Arabidopsis. The Plant cell 19: 473–84.

Koyama T, Sato F, Ohme-Takagi M. 2017. Roles of miR319 and TCP transcription factors in leaf development. Plant Physiology 175: pp.00732.2017.

Lifschitz E, Eviatar T, Rozman A, Shalit A, Goldshmidt A, Amsellem Z, Alvarez JP, Eshed Y. 2006. The tomato *FT* ortholog triggers systemic signals that regulate growth and flowering and substitute for diverse environmental stimuli. Proc Natl Acad Sci USA 103: 6398–6403.

Lintilhac PM. 2014. The problem of morphogenesis : unscripted biophysical control systems in plants. : 25–36.

Maltnan DS, Jenkins J a LB-PBSR 20. 1962. A morphogenetic study of Lanceolate, a leaf shape mutant in tomato. Am. J. Bot. 49: 504–514.

Mathan DS, Jenkins JA. 1962. A Morphogenetic Study of Lanceolate, A Leaf-Shape Mutant in the Tomato. American Journal of Botany 49: 504–514.

Maugarny-Calès A, Laufs P. 2018. Getting leaves into shape: a molecular, cellular, environmental and evolutionary view. Development 145: 1–16.

Menda N, Semel Y, Peled D, Eshed Y, Zamir D. 2004. In silico screening of a saturated mutation library of tomato. The Plant journal□: for cell and molecular biology 38: 861–872.

Munjal A, Philippe J, Munro E, Lecuit T. 2015. A self-organized biomechanical network drives shape changes during tissue morphogenesis.

Nath U, Crawford BCW, Carpenter R, Coen E. 2003. Genetic Control of Surface Curvature. Science 299: 1404–1408.

Naz AA, Raman S, Martinez CC, Sinha NR, Schmitz G, Theres K. 2013. Trifoliate encodes an MYB transcription factor that modulates leaf and shoot architecture in tomato. Proceedings of the National Academy of Sciences of the United States of America 110: 2401–6.

Nir I, Shohat H, Panizel I, Olszewski NE, Aharoni A, Weiss D. 2017. The Tomato DELLA Protein PROCERA Acts in Guard Cells to Promote Stomatal Closure. The Plant Cell: tpc.00542.2017.

Ori N, Cohen AR, Etzioni A, Brand A, Yanai O, Shleizer S, Menda N, Amsellem Z, Efroni I, Pekker I, et al. 2007. Regulation of LANCEOLATE by miR319 is required for compound-leaf development in tomato. Nature genetics 39: 787–791.

Palatnik JF, Allen E, Wu X, Schommer C, Schwab R, Carrington JC, Weigel D. 2003. Control of leaf morphogenesis by microRNAs. Nature 425: 257–263.

Pałubicki W, Kokosza A, Burian A. 2019. Formal description of plant morphogenesis. Journal of Experimental Botany 70: 3601–3613.

Petit F, Sears KE, Ahituv N. 2017. Limb development : a paradigm of gene regulation. Nature 18: 245–258.

Poethig RS. 1997. Leaf morphogenesis in flowering plants. The Plant cell 9: 1077–1087.

Raman S, Greb T, Peaucelle A, Blein T, Laufs P, Theres K. 2008. Interplay of miR164, CUP-SHAPED COTYLEDON genes and LATERAL SUPPRESSOR controls axillary meristem formation in Arabidopsis thaliana. Plant Journal 55: 65–76.

Rast-somssich MI, Broholm S, Jenkins H, Canales C, Vlad D, Kwantes M, Bilsborough G, Ioio R Dello, Ewing RM, Laufs P, et al. 2015. Alternate wiring of a KNOXI genetic network underlies differences in leaf development of A. thaliana and C. hirsuta Madlen. Genes & Development 29(22): 2391–2404.

Rodriguez RE, Debernardi JM, Palatnik JF. 2014. Morphogenesis of simple leaves: Regulation of leaf size and shape. Wiley Interdisciplinary Reviews: Developmental Biology 3: 41–57.

Sakamoto T, Kamiya N, Ueguehi-Tanaka M, Iwahori S, Matsuoka M. 2001. KNOX homeodomain protein directly suppresses the expression of a gibberellin biosynthetic gene in the tobacco shoot apical meristem. Genes and Development 15: 581–590.

Schommer C, Debernardi JM, Bresso EG, Rodriguez RE, Palatnik JF. 2014. Repression of cell proliferation by miR319-regulated TCP4. Molecular Plant 7: 1533–1544.

Scofield S, Dewitte W, Nieuwland J, Murray JAH. 2013. The Arabidopsis homeobox gene SHOOT MERISTEMLESS has cellular and meristem-organisational roles with differential requirements for cytokinin and CYCD3 activity. The Plant Journal 75: 53–66.

Shani E, Ben-Gera H, Shleizer-Burko S, Burko Y, Weiss D, Ori N. 2010. Cytokinin regulates compound leaf development in tomato. The Plant cell 22: 3206–3217.

Shani E, Burko Y, Ben-Yaakov L, Berger Y, Amsellem Z, Goldshmidt A, Sharon E, Ori N. 2009. Stage-specific regulation of Solanum lycopersicum leaf maturation by class 1 KNOTTED1-LIKE HOMEOBOX proteins. The Plant cell 21: 3078–3092.

Shleizer-Burko S, Burko Y, Ben-Herzel O, Ori N. 2011. Dynamic growth program regulated by LANCEOLATE enables flexible leaf patterning. Development (Cambridge, England) 138: 695–704.

Shwartz I, Levy M, Ori N, Bar M. 2016. Hormones in tomato leaf development. Developmental Biology 419: 132–142.

Silva GFF, Silva EM, Correa JPO, Vicente MH, Jiang N, Notini MM, Junior AC, De Jesus FA, Castilho P, Carrera E, et al. 2019. Tomato floral induction and flower development are orchestrated by the interplay between gibberellin and two unrelated microRNA-controlled modules. New Phytologist 221: 1328–1344.

Šimura J, Antoniadi I, Široká J, Tarkowská D, Strnad M, Ljung K, Novák O. 2018. Plant Hormonomics : Multiple Phytohormone Profiling by. Plant Physiology 177:476–489.

Speybroeck L Van, Waele D De, Vijver G Van De. 2006. Theories in Early Embryology. Ann. N.Y. Acad. Sci. 981: 7–49.

Steiner E, Livne S, Kobinson-Katz T, Tal L, Pri-Tal O, Mosquna A, Tarkowská D, Mueller B, Tarkowski P, Weiss D. 2016. The putative O-linked N-acetylglucosamine transferase SPINDLY inhibits class I TCP proteolysis to promote sensitivity to cytokinin. Plant Physiology 171: 1485–1494.

Sutherland A, Keller R, Lesko A. 2020. Seminars in Cell & Developmental Biology Convergent extension in mammalian morphogenesis. Seminars in Cell and Developmental Biology 100: 199–211.

Theophrastus. 1916. Enquiry into Plants 350–285 BC, trans Hort A. Harvard Univ Press, Cambridge, MA.

Weiss D, Ori N. 2007. Mechanisms of cross talk between gibberellin and other hormones. Plant physiology 144: 1240–6.

Yanai O, Shani E, Dolezal K, Tarkowski P, Sablowski R, Sandberg G, Samach A, Ori N. 2005. Arabidopsis KNOXI proteins activate cytokinin biosynthesis. Current biology□: CB 15: 1566–71.

Yanai O, Shani E, Russ D, Ori N. 2011. Gibberellin partly mediates LANCEOLATE activity in tomato. The Plant journal□: for cell and molecular biology 68: 571–582.

Zürcher E, Tavor-Deslex D, Lituiev D, Enkerli K, Tarr PT, Müller B. 2013. A robust and sensitive synthetic sensor to monitor the transcriptional output of the cytokinin signaling network in planta. Plant physiology 161: 1066–75.

