## Supplementary material for "Coordinating the morphogenesis-differentiation balance by tweaking the cytokinin-gibberellin equilibrium": description of supplemetal datasets

Supplemental dataset 1- Lists of upregulated DEGs in all genotypes- excel file.

Tab 1: Genes upregulated in *clausa.*

Tab 2: Genes downregulated in *clausa*.

Tab 3: Genes Upregulated in *La2/+.*

Tab 4: Genes Downregulated in *La2/+.*

Tab 5: Genes Upregulated in *la6*.

Tab 6: Genes Downregulated in *la6*.

Tab 7: Genes upregulated in *pFIL>>miR319.*

Tab 8: Genes Downregulated in *pFIL>>miR319*.

Tab 9: Genes upregulated in *pFIL>>TKN2*.

Tab 10: Genes downregulated in *pFIL>>TKN2*.

Tab 11: Genes upregulated in *pFIL>>TKN2-SRDX*.

Tab 12: Genes downregulated in *pFIL>>TKN2-SRDX*.

Tab 13: Comparison of all groups.

Tab 14: pairwise comparisons between groups.

Tab 15: Representation Factor (RF) of comparisons between groups.

Supplemental dataset 2- Differential KEGG pathways in all genotypes- excel file.

Tab 1: KEGG pathways down in *La2/+*.

Tab 2: KEGG pathways up in *La2/+*.

Tab 3: KEGG pathways down in *pFIL>>miR319*.

Tab 4: KEGG pathways up in *pFIL>>miR319*.

Tab 5: KEGG pathways down in *la6*.

Tab 6: KEGG pathways up in *la6*.

Tab 7: KEGG pathways down in *clausa*.

Tab 8: KEGG pathways up in *clausa.*

Tab 9: KEGG pathways down in *pFIL>>TKN2*.

Tab 10: KEGG pathways up in *pFIL>>TKN2*.

Tab 11: KEGG pathways down in *pFIL>>TKN2*-*SRDX*.

Tab 12: KEGG pathways up in *pFIL>>TKN2*-*SRDX*.

Tab 13: KEGG pathways common to several genotypes.

Tab 14: KEGG pathways common different genotypes with venn diagrams.

Supplemental dataset 3- morphogenetic and diff genes from Ichihashi et al., 2014- excel file.

Tab 1: "Morphogenetic" genes from Ichihashi et. al., 2014, and their behavior in selected genotypes.

Tab 2: "Differentiative" genes from Ichihashi et. al., 2014, and their behavior in selected genotypes.

Supplemental dataset 4 (or figure X)- Representation factor and analysis of the overlap of "morphogenetic" and "differentiative" genes from Ichihashi et al., 2014, with CLAU and LA genotypes. Word doc.

Supplemental dataset 5 – KNOX putative targets?
