## Supplementary material for "Coordinating the morphogenesis-differentiation balance by tweaking the cytokinin-gibberellin equilibrium": Figures

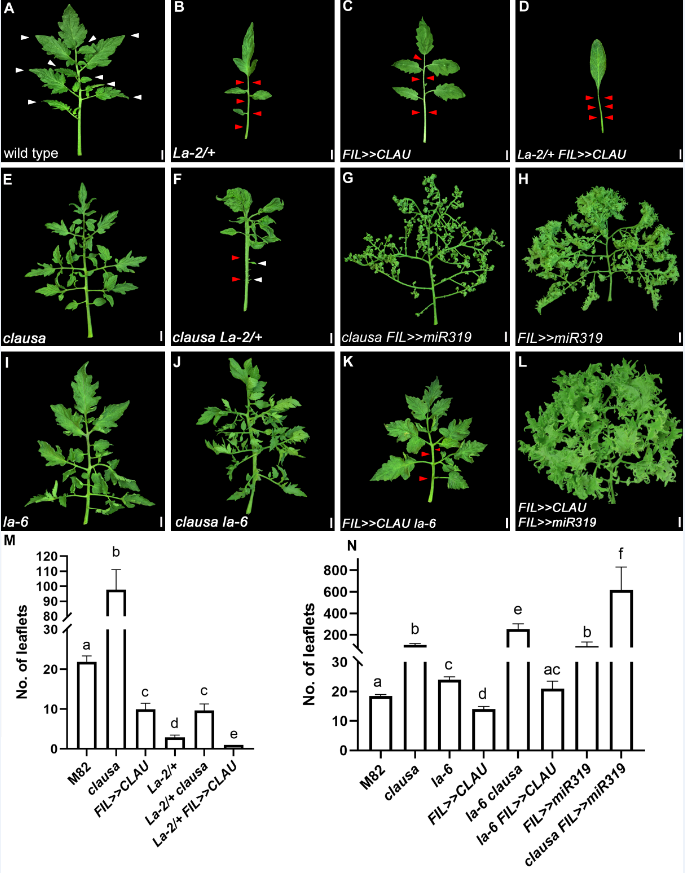
**Figure 1**

***CLAU* and *LA* function in parallel pathways.**

**(A-L)** Genetic interactions between genotypes with altered *CLAU* and *LA* expression levels. *La-2/+*: a semi-dominant LA allele with increased and precocious expression due to miR319 resistance. *La-6*, *FIL>>MIR319*: *LA* null or *LA* downregulation, respectively. *clausa*: *CLAU* null. *FIL>>CLAU*: *CLAU* upregulation. All leaves depicted are fully expanded fifth leaves. Bars = 1 cm.

White and red arrowheads represent primary leaflets and missing primary and intercalary leaflets, respectively.

**(M-N)** Quantification of leaf complexity in genotypes with altered *CLAU* and *LA* expression levels. Graphs represent mean ± SE of six independent biological repeats. Statistical significance of differences was examined in a one-way ANOVA, p<0.0001. Different letters indicate significant differences between samples in an unpaired two-tailed t-test with Welch's correction.

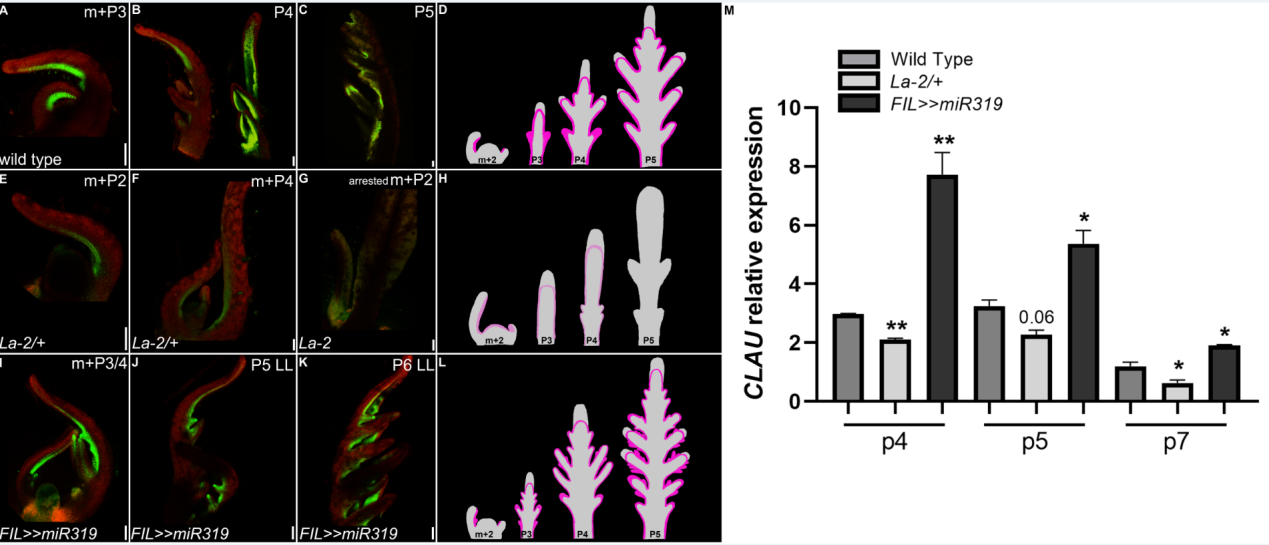

**Figure 2**

***LA* determines the window for *CLAU* expression**

**(A-L)** Expression of the *CLAUSA* promoter *CLAU::nYFP* in altered *LA* genotypes. **(A-C)** Expression of *CLAU::nYFP* in different developmental stages of WT; **(E-G)** Expression of *CLAU::nYFP* in different developmental stages of *La-2/+* (*LA* upregulation); **(I-K)** Expression of *CLAU::nYFP* in different developmental stages of *FIL>>MIR319* (Reduced *LA* activity). The pattern of *CLAU::nYFP* expression was detected by a confocal laser scanning microscope (CLSMmodel SP8; Leica), with the solid-state laser set at 514 nm excitation/ 530 nm emission. Chlorophyll expression was detected at 488nm excitation/ 700nm emission. Bars = 100 um

**(D, H, L)**: Cartoon summarizing the expression of *pCLAU::nYFP* throughout development in each genotype.

**(M)** Expression levels of *CLAU* in altered *LA* genotypes was determined in successive leaf developmental stages using RT-qPCR. Graphs represent mean ± SE of three independent biological repeats. Asterisks indicate significant differences from *CLAU* expression in WT in an unpaired two-tailed t-test, p≤0.0387.

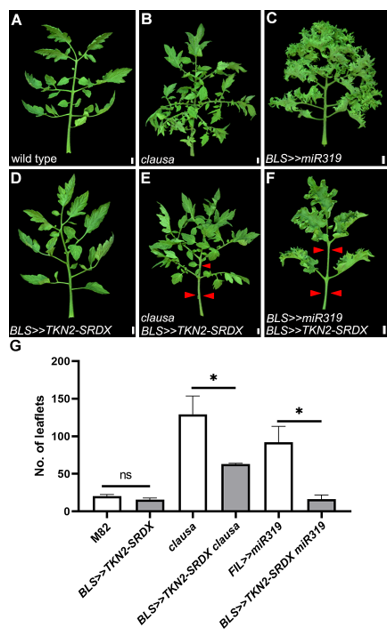

**Figure 3**

**Inhibiting TKN2 targets significantly reduces morphogenesis in *CLAU-* and *LA-*deficient backgrounds**

**(A-F)** Overexpression of *TKN2-SRDX* in the background of *CLAU* and *LA* deficiency. All leaves depicted are fully expanded fifth leaves. Bars = 1 cm.

Red arrowheads represent primary leaflets and missing primary and intercalary leaflets.

**(G)** Quantification of leaf complexity upon overexpression of TKN2-SRDX in the background of *CLAU* and *LA* deficiency. Graphs represent mean ± SE of at least three independent biological repeats. Asterisks indicate significant differences from the background genotype (without TKN2-SRDX overexpression) in an unpaired two-tailed t-test with Welch's correction, p≤0.0335.

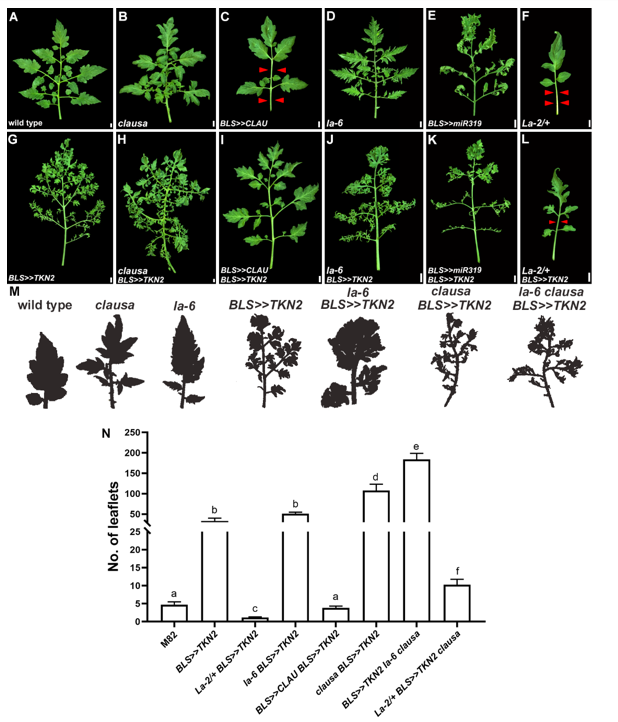

**Figure 4**

**TKN2 promotes morphogenetic activity in altered *CLAU* and *LA* backgrounds.**

**(A-L)** Overexpression of *TKN2* in the background of genotypes with altered *CLAU* and *LA* expression levels. All leaves depicted are fully expanded fifth leaves. Bars = 1 cm.

Red arrowheads represent missing primary and intercalary leaflets.

**(M)** Shaded cartoon of the terminal leaflet of the indicated genotypes.

**(N)** Quantification of leaf complexity upon overexpression of TKN2 in genotypes with altered *CLAU* and *LA* expression levels. Graphs represent mean ± SE of at least three independent biological repeats. Letters indicate significant differences between samples in a one-way ANOVA with a Tukey post-hoc test, p<0.027.

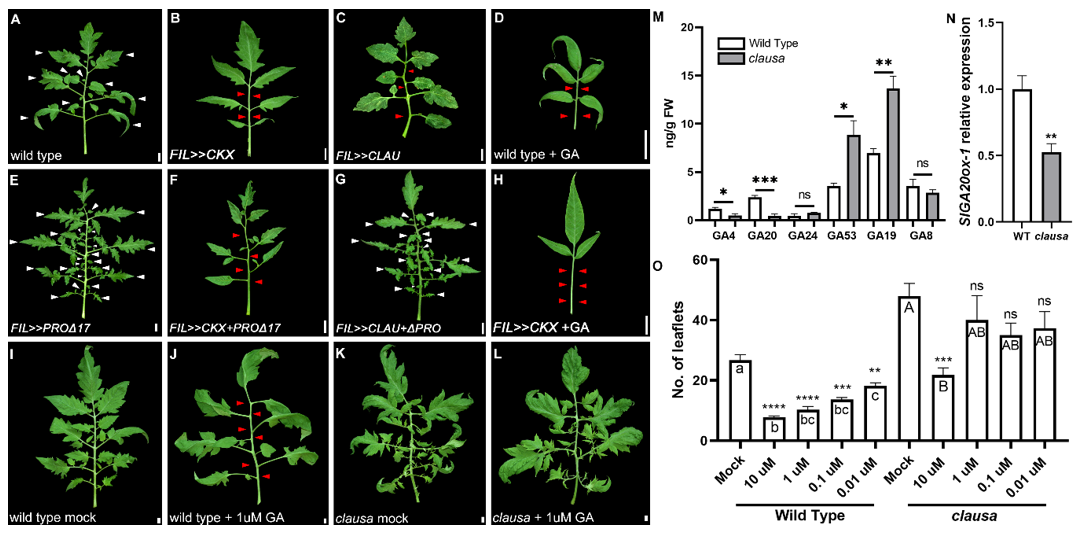
**Figure 5**

***clausa* has an altered GA profile and reduced sensitivity to GA treatment**

**(A-D)** Phenotypes of leaves with reduced CK (*FIL>>CKX*) increased CLAU (*FIL>>CLAU*), or that received exogenous GA. **(E-H)** Phenotypes of leaves with reduced GA response (*FIL>>∆PRO*), reduced CK and reduced GA response (*FIL>>CKX*+*∆*PRO), or increased CLAU and reduced GA response (*FIL>>CLAU*+ *∆*PRO). **(I-L)** Effect of GA treatment on WT and *clausa*. All leaves depicted are fully expanded fifth leaves. Bars = 1 cm.

White and red arrowheads represent primary leaflets and missing primary and intercalary leaflets, respectively.

**(M)** Quantification of GAs in WT and *clausa*. Asterisks indicate significant differences between WT and *clausa* for each GA in an unpaired two-tailed t-test, *p≤0.05, **p≤0.01, ***p≤0.001.

**(N)** Expression of *SlGA20ox-1,* the enzyme that converts GA19 to GA20*,* in WT and *clausa*, was determined in young shoots (m+6) of 2-week-old plants using RT-qPCR. Graphs represent mean ± SE of five independent biological repeats. Asterisks indicate significant differences in an unpaired two-tailed t-test, p=0.0078.

**(O)** Quantification of leaf complexity following GA treatment in WT and *clausa*. Graphs represent mean ± SE of at least three independent biological repeats. Asterisks and letters indicate significant differences between samples (asterisks= GA treatments compared to Mock in each genotype) in a Welch's one-way ANOVA with a Dunnett post-hoc test, letters: p<0.0097; **p<0.01, ***p<0.001, ****p<0.0001.

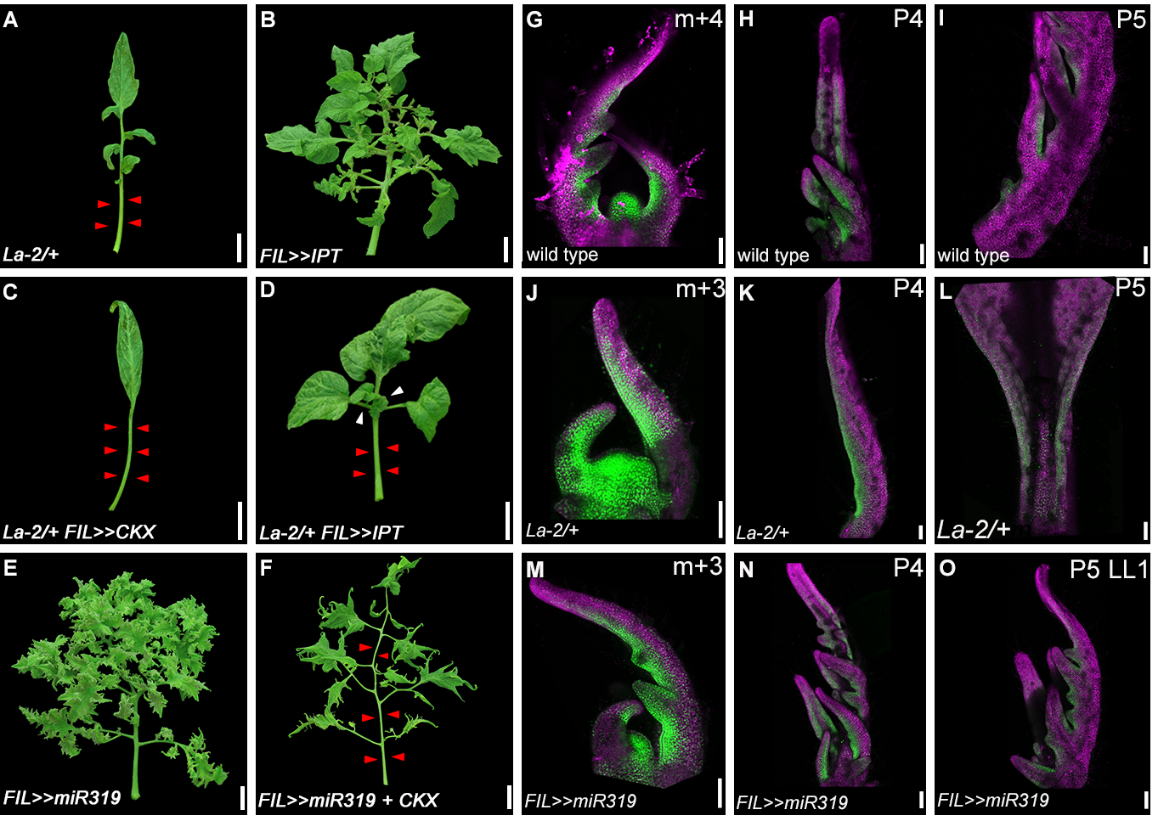

**Figure 6**

***LA* reduces leaf margin sensitivity to CK**

(A-F) Phenotypes of leaves with increased CK (*FIL>>IPT*) or decreased CK (*FIL>>CKX*) levels, and increased *LA* activity (*La-2/+*) or reduced *LA* activity (*FIL>>miR319).* All leaves depicted are fully expanded fifth leaves. Bars = 2 cm.

White and red arrowheads represent primary leaflets and missing primary and intercalary leaflets, respectively.

(G-O) Confocal micrographs of *TCSv2::3XVENUS* in successive developmental stages in indicated genotypes. *TCSv2* driven signals are reduced in *La2/+* in the fifth plastochron. The pattern of VENUS expression was detected by a confocal laser scanning microscope (CLSMmodel SP8; Leica), with the solid-state laser set at 514 nm for excitation and 530 nm for emission. Chlorophyll emission was detected at 488nm excitation/ 700nm emission. Bars = 100 um.

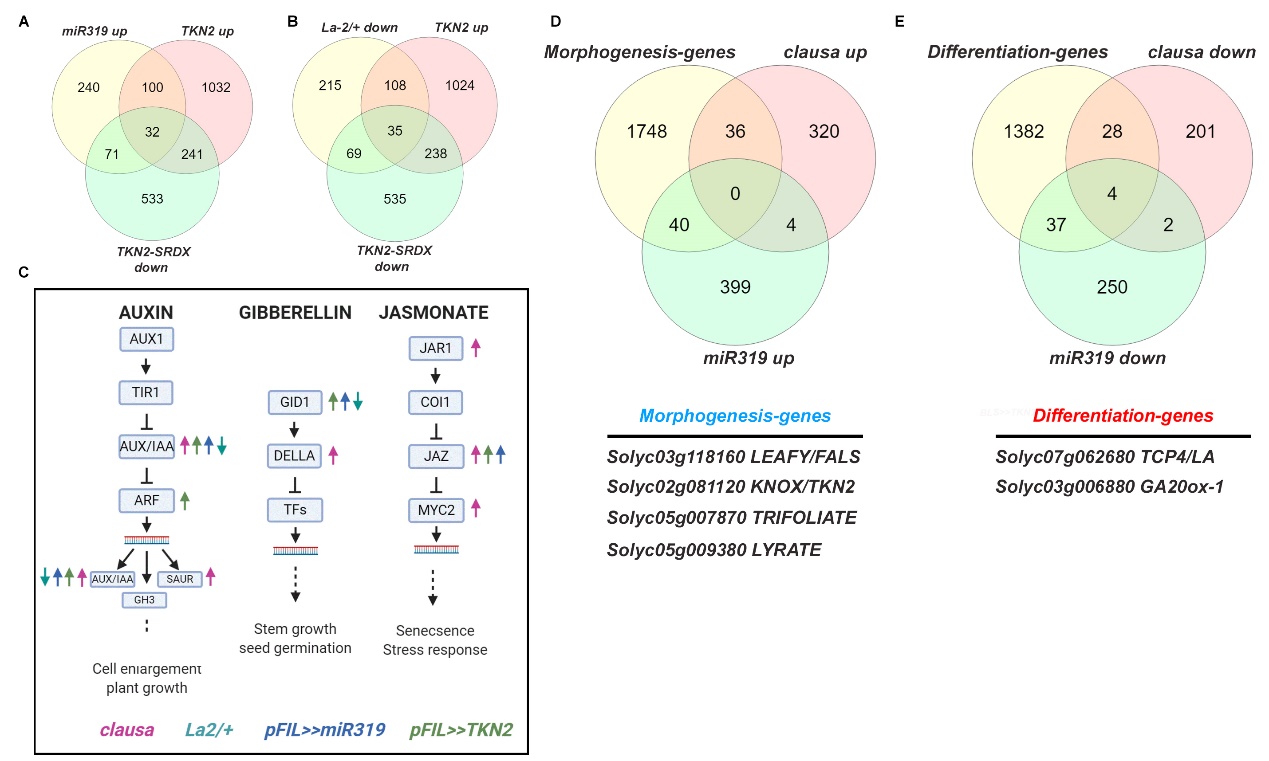

**Figure 7**

***Global expression analyses of LA, CLAU and TKN2***

(A-B) Venn diagrams depicting co-regulated genes in LA and TKN2 genotypes. Roughly a third to half of the genes upregulated upon LA activity downregulation (*miR319* overexpression), or downregulated upon LA activity upregulation (*La-2/+*), are co-regulated upon upregulation of TKN2 activity (overexpression of *TKN2*) and/or downregulation of the activity of TKN2 targets (overexpression of *TKN2-SRDX*).

(C) Hormone signaling pathways co-regulated in the different genotypes. Arrow color corresponds with genotypes provided below, arrow direction indicates up or down regulation.

(D-E) Venn diagrams depicting co-regulated morphogenesis and differentiation genes upon decrease of CLAU activity (*clausa*) and LA activity (*miR319* overexpression). Genes were curated from (Ichihashi *et al.*, 2014).

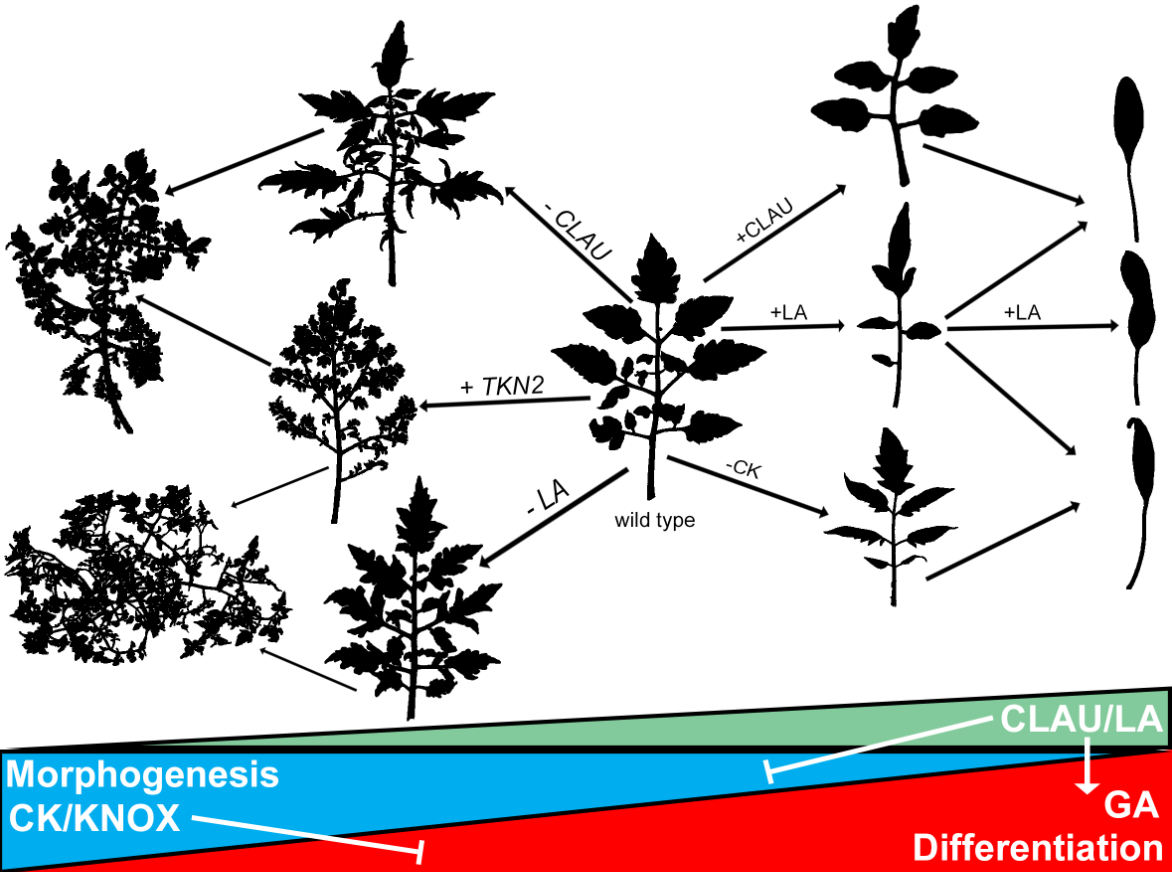

**Figure 8**

**Model for the role of *CLAU*, *LA and TKN2* and GA-CK balance in leaf development**

CK promotes morphogenesis, and GA promotes differentiation. Increasing CLAU or LA levels leads to increased GA sensitivity and decreased CK sensitivity, pulls the leaf developmental program towards differentiation, and results in simplified leaf forms. Decreasing CLAU or LA levels leads to increased CK sensitivity and decreased GA sensitivity, pulls the leaf developmental program towards morphogenesis and results in elaborate leaf forms. LA acts in a dose-dependent manner. While only 1 copy of the dominant LA allele in *La-2/+* gain-of-function mutants produces some leaflets, adding additional copy of LA allele in the *La-2* homozygous plants produces a completely simple leaf. The additional LA copy can be replaced by increasing CLAU levels, by reducing CK levels or by increasing GA levels (Mathan & Jenkins, 1962). Reducing CLAU or LA activity or increasing TKN2 activity leads to increase in leaf complexity, which is further enhanced when combining two of them.

**Supplemental Figures**

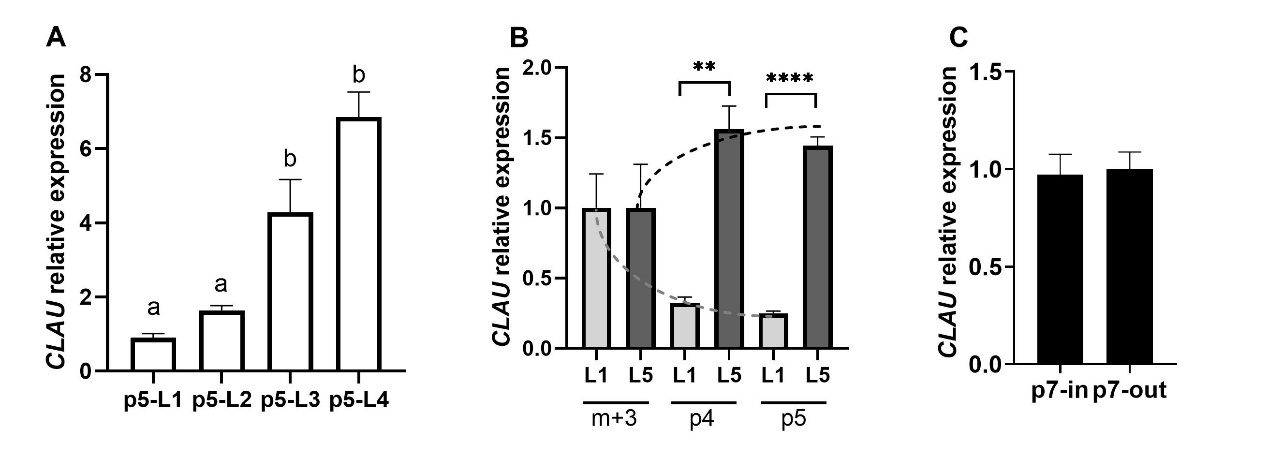

**Figure S1**

***CLAU* expression in successive developmental stages of different leaves**

Expression level of *CLAU* was determined using RT-qPCR. **(A)** *CLAU* expression in the fifth plastochron in leaves 1 through 4. Leaves 1 and 2, which have reduced complexity, also have reduced CLAU expression. Graph represents mean ± SE of three independent biological repeats. Letters indicate significant differences in a one-way ANOVA with a Tukey post-hoc test, p≤0.037. **(B)** *CLAU* expression in successive leaf developmental stages, comparing the first and fifth leaves. Dashed line indicates expression trend. Graph represents mean ± SE of three independent biological repeats. Asterisks indicate significant differences in an unpaired, two-tailed t-test, p≤0.0019. **(C)** *CLAU* expression in the inner and outer regions of the fifth leaf at the seventh plastochron stage. Graph represents mean ± SE of three independent biological repeats. No significant differences.

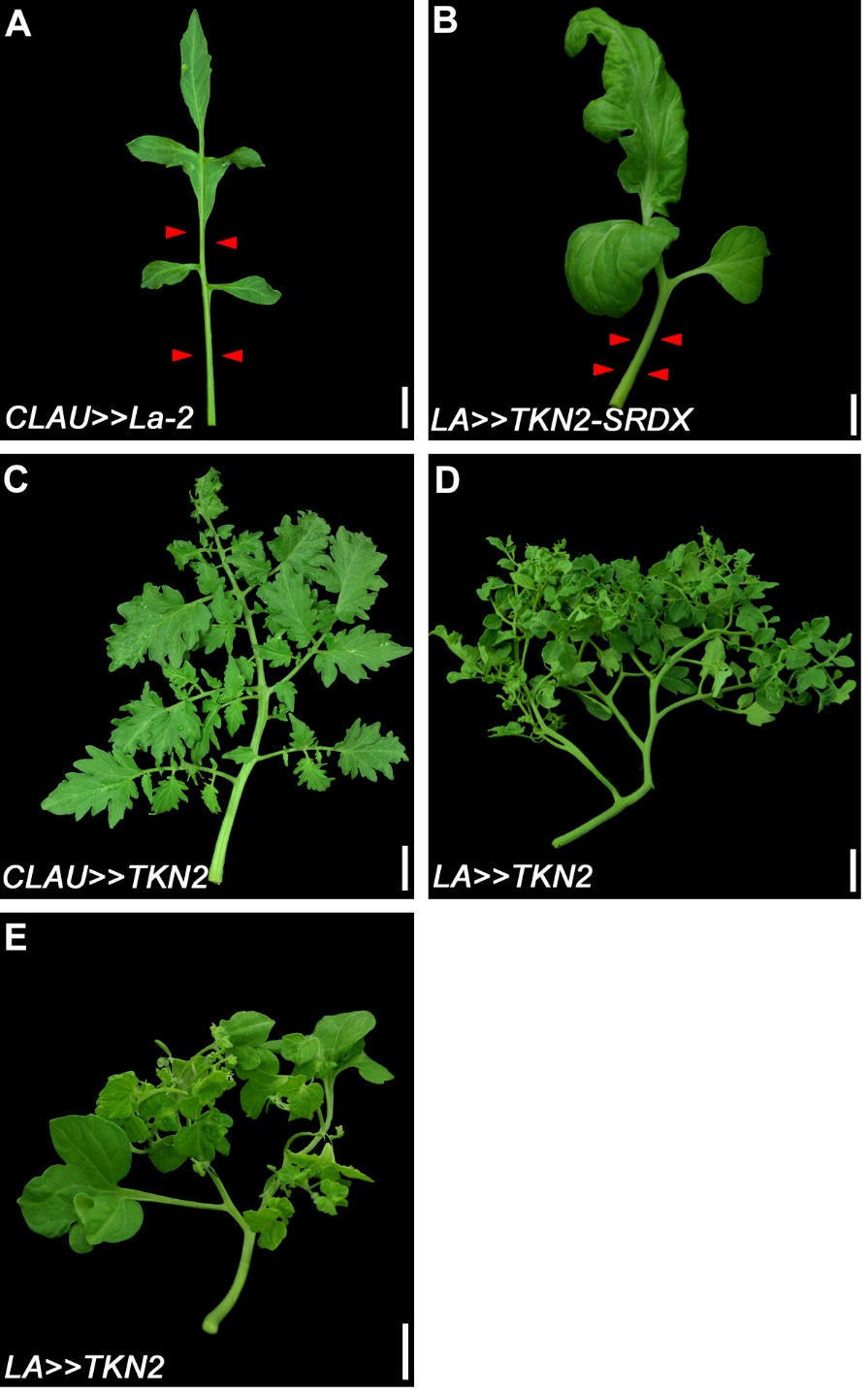

**Figure S2**

**LA and CLAU expression domains and activity and TKN2**

**(A-E)** Phenotypes of leaves of the indicated genotypes*.* All leaves depicted are fully expanded fifth leaves. Bars = 2 cm.

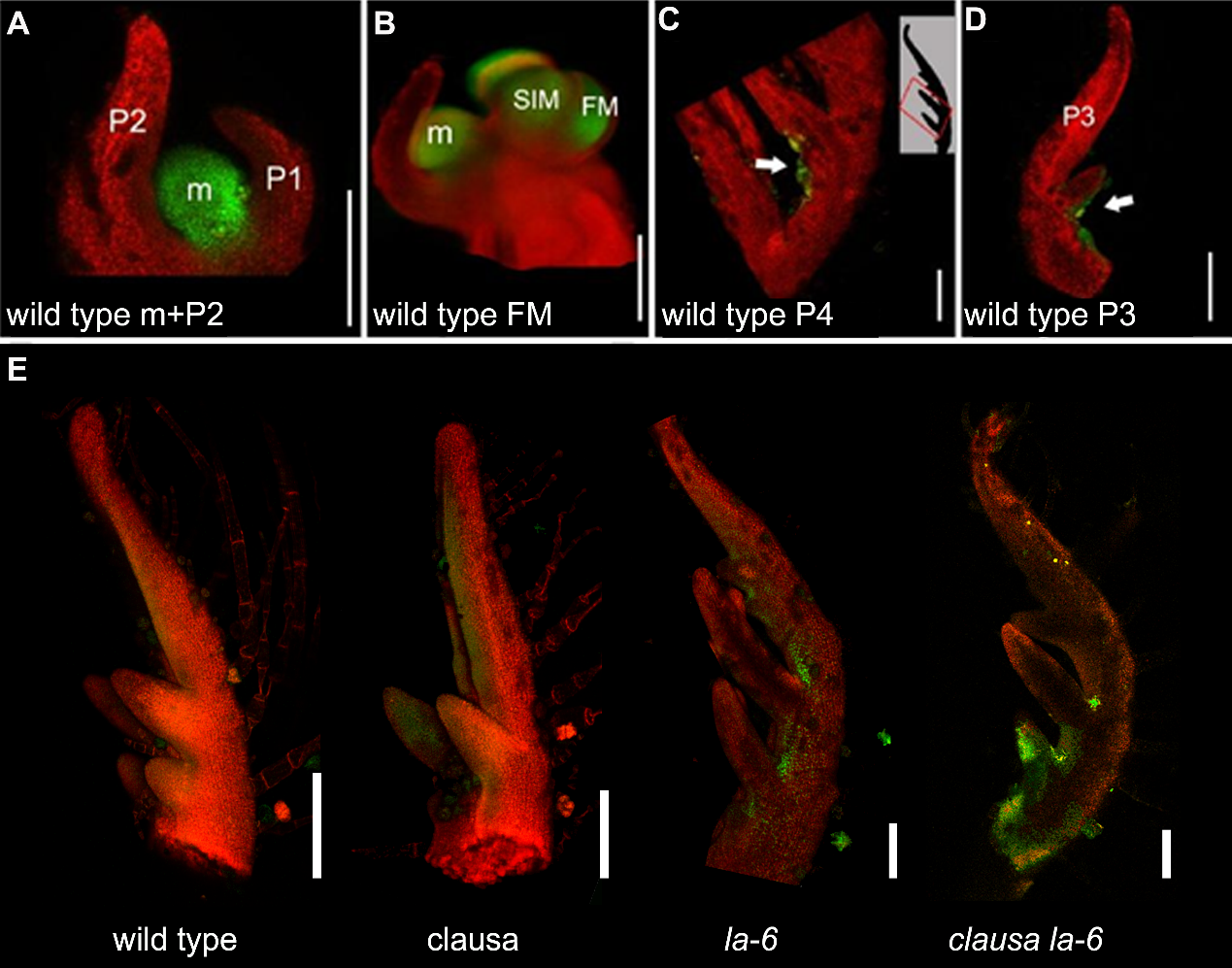

**Figure S3**

**TKN2 is expressed at the leaf margin in *CLAU* and *LA* deficient backgrounds**

**(A-D)** Expression pattern of the TKN2 promoter fused to YFP in WT M82 plants. **(A)** Vegetative meristem (m) and two youngest leaf primordia (P1, P2). **(B)** Floral meristem (FM), sympodial inflorescence meristem (SIM). **(C-D)** Weak expression is observed in P3/P4 leaf primordia (area of the leaf primordium depicted in C is indicated in the inset). **(E)** Confocal micrographs of *pTKN::nYFP* in the fourth plastochron (P4) in indicated genotypes. TKN2 promoter activation increases in the leaf margin in *CLAU* and *LA* deficient backgrounds.

The pattern of YFP expression was detected by a confocal laser scanning microscope (CLSMmodel SP8; Leica), with the solid-state laser set at 514 nm for excitation and 530 nm for emission. Chlorophyll emission was detected at 488nm excitation/ 700nm emission. Bars = 200 um.

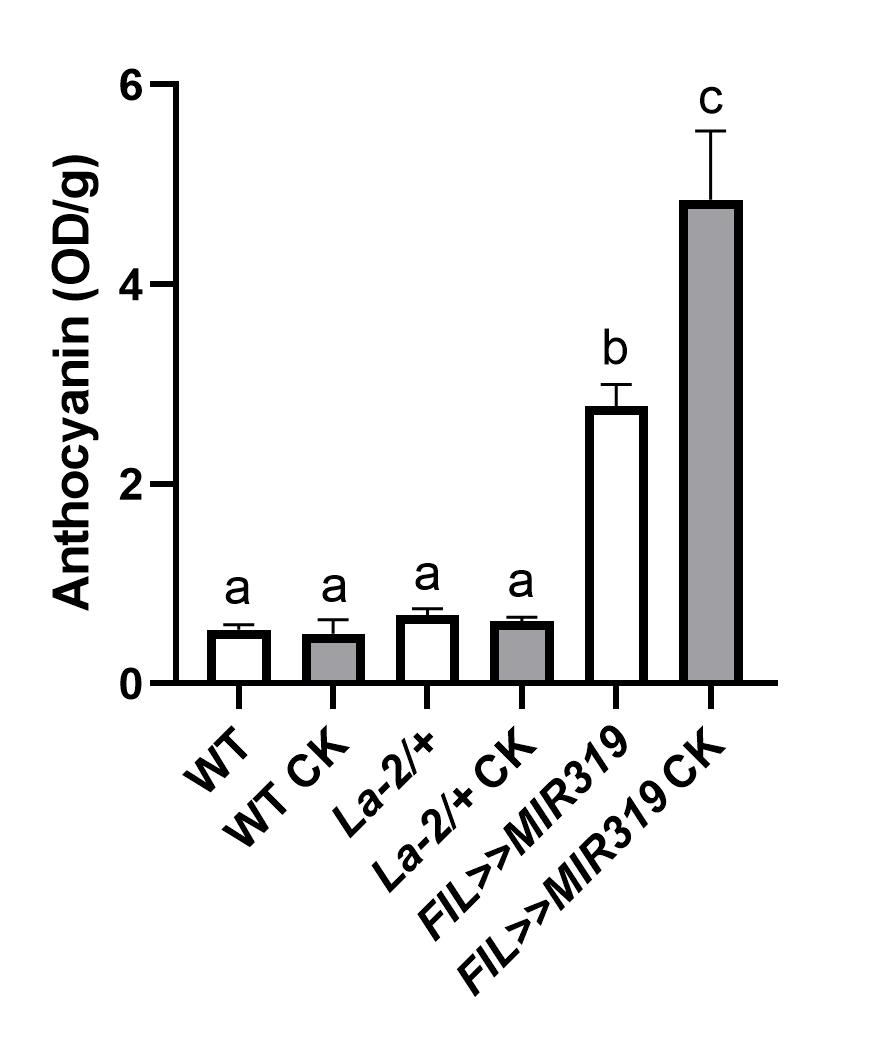

**Figure S4**

***LA* deficiency enhances CK sensitivity to anthocayanin accumulation.**

Anthocyanin content in WT and altered *LA* genotypes with or without CK treatment was determined by measuring optical density following methanolic extraction. Graph represents mean ± SE of five independent biological repeats. Letters indicate significant differences in an unpaired, two-tailed t-test with Welch's correction, p≤0.038.

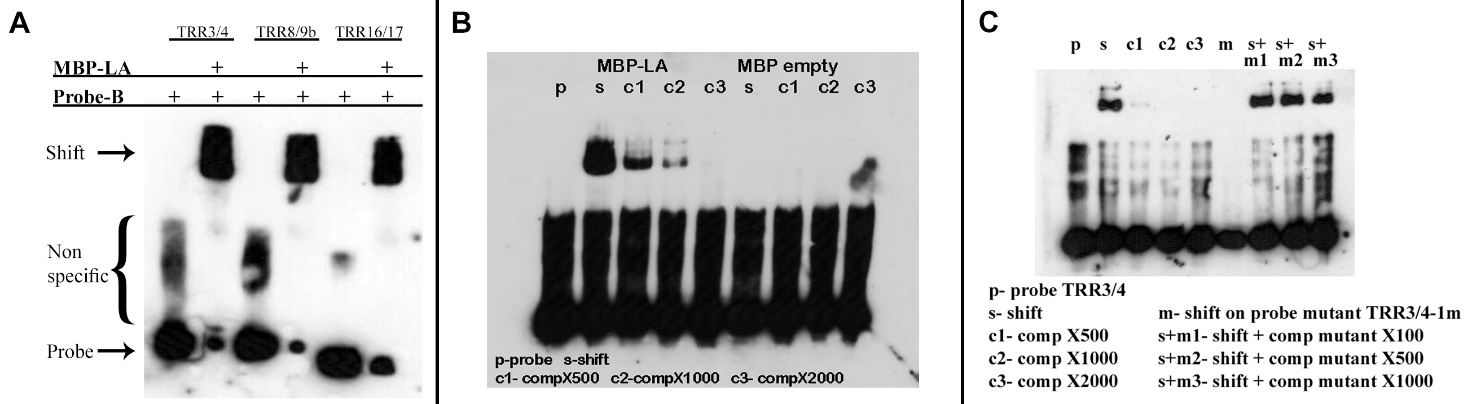

**Figure S5**

**LA binds *in vitro* to the promoters of TRRs.**

**A** LA binds *in vitro* to the promoters of TRRs. Recombinant MBP-LA caused a shift in the PAGE migration of biotinylated promoter derived probes (Probe-B) of the indicated TRRs.

**B** Specific binding of *LA* to the TRR3/4 promoter. Recombinant MBP-LA caused a shift (**s**) in the PAGE migration of the biotinylated promoter derived probe (**p**) of TRR3/4. A non-biotinylated probe was able to compete with the biotinylated probe in a quantity depended manner (**c1-c3**). MBP (**MBP empty**) was used as a control protein and does not cause a probe shift in PAGE.

**C** A biotinylated probe with a mutation in the binding site (**m**) does not shift in the presence of MBP-LA, and cannot compete with the WT probe (**s+m1-m3**).

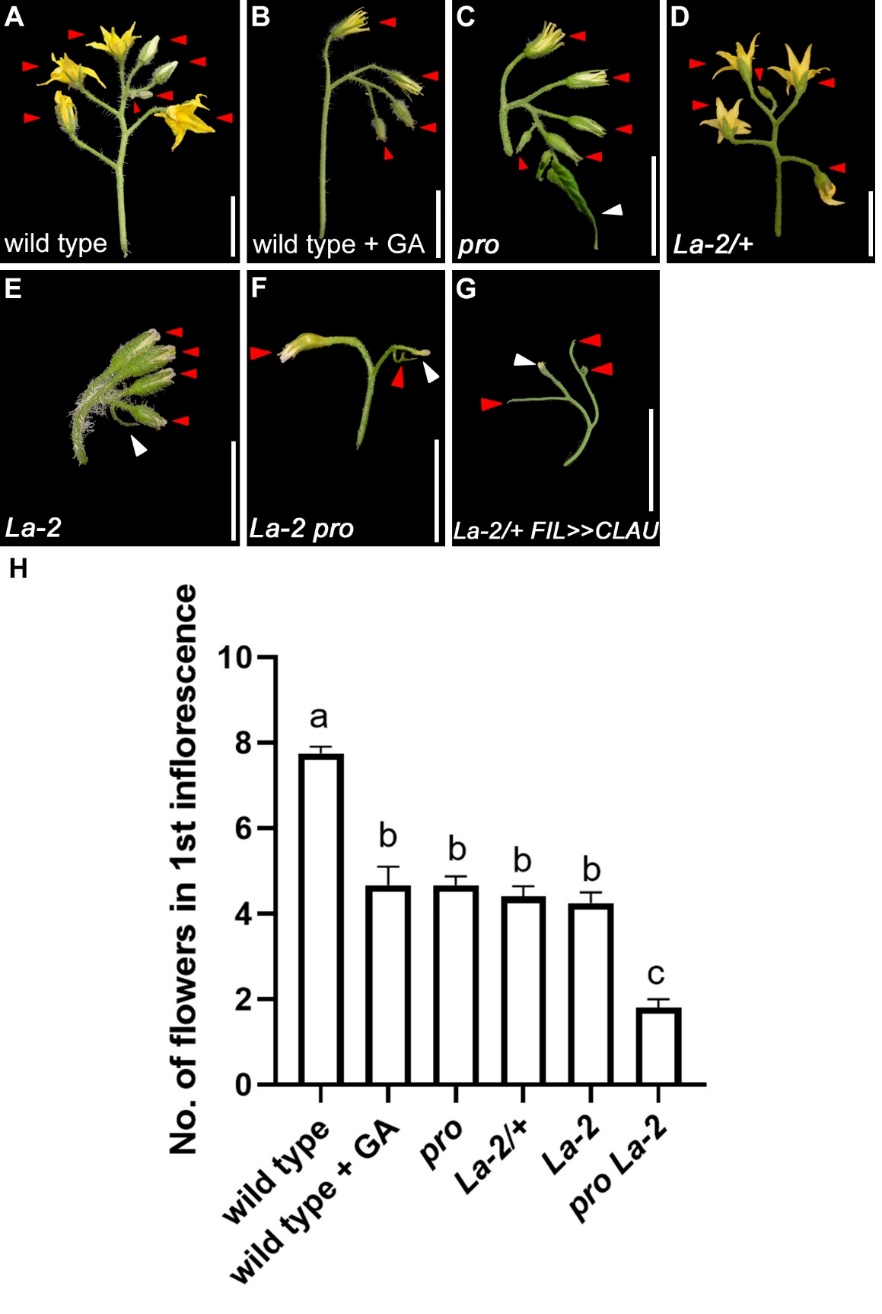

**Figure S6**

***CLAU* and *LA* function in parallel pathways in additional developmental processes.**

**(A-H)** Genetic interactions between genotypes with altered *CLAU* and *LA* expression levels and GA levels or response. *La-2/+*: a semi-dominant LA allele with increased and precocious expression due to miR319 resistance. *FIL>>CLAU*: *CLAU* upregulation. Increased GA levels (GA application) or response: in *procera (*pro) loss-of-function. Bars = 2 cm.

Red and white arrowheads represent flowers and leaf-like structures, respectively.

**(I)** Quantification of inflorescence complexity in the indicated genotypes and treatments. Graphs represent mean ± SE of at least 4 independent biological repeats. Letters indicate significant differences between samples in a one-way ANOVA with a Tukey post-hoc test, p<0.0009.

**Supplemental Table 1**

**Primers used in this work**

| **Name** | **5'-3' seq** | **Use** |
| --- | --- | --- |
| pTKN2-R | CAC CCT GTC TCT AAT GAT CTC CAT CCC TA | pTKN2 cloning |
| pTKN2 -L | CAT TAT TCT CTC ACA CAC TTT CTT CTT | pTKN2 cloning |
| EXP RT-F | TGGGTGTGCCTTTCTGAATG | qRT-PCR |
| EXP RT-R | GCTAAGAACGCTGGACCTAATG | qRT-PCR |
| Clau F | CCTCTCACAACAAGCAATGAACTT | qRT-PCR |
| Clau R | AGGACGATGCAATGAGAGAGAC | qRT-PCR |
| GA2ox F | CACCATGGCTATTGATTGTATGATCACAAATG | qRT-PCR |
| GA2ox R | TTAAGCTTGTGTAGTAGTGTTGTGTTG | qRT-PCR |
| TRR16/17-1 F | AAGAATGGGAAGCAGCTAGATTTGGTCCCCATTATTAAGATTTAGCAAGA | EMSA probe generation |
| TRR16/17-1 R | TCTTGCTAAATCTTAATAATGGGGACCAAATCTAGCTGCTTCCCATTCTT | EMSA probe generation |
| TRR8/9-1 F | CTATTAGATTACACTTGTAAAACGTGTGGTCCAACATAGAGAAATGGTAATCTTTTTC | EMSA probe generation |
| TRR8/9-1 R | AAAAAGATTACCATTTCTCTATTGTTGGACCACACGTTTTACAAGTGTAATCTAATAGC | EMSA probe generation |
| TRR3/4-1 F | AAAAAAGGTAAATATATAAATTATGTAGGACCACAAAAAGATTTTGTAAAGATTAGACAA | EMSA probe generation |
| TRR3/4-1 R | TTGTCTAATCTTTACAAAATCTTTTTGTGGTCCTACATAATTTATATATTTACCTTTTTT | EMSA probe generation |
| TRR3/4-1m F | AAAAAAGGTAAATATATAAATTATGTAGGAACTCAAAAAGATTTTGTAAAGATTAGACAA | EMSA probe generation |
| TRR3/4-1m R | TTGTCTAATCTTTACAAAATCTTTTTGAGTTCCTACATAATTTATATATTTACCTTTTTT | EMSA probe generation |
