## Supplementary material for "Coordinating the morphogenesis-differentiation balance by tweaking the cytokinin-gibberellin equilibrium": Supp dataset4- morphogenesis and differentiation

Morphogenesis and differentiation genes were identified from a published dataset (Ichihashi et al., 2014).

1824 genes fit the condition of m+P3 exp> P4 exp> P5 exp> P6 exp and are considered "Morphogenetic" genes.

1451 genes fit the condition of m+P3 exp< P4 exp< P5 exp< P6 exp and are considered "differentiative" genes.

"Morphogenetic" genes, as a group, are significantly enriched in the upregulated group in *clausa*, *pFIL>>miR319*, and *pFIL>>TKN2*, and in the downregulated group in *La2/+* and *pFIL>>TKN2-SRDX*. They are also significantly depleted in the downregulated group in *clausa* and *pFIL>>TKN2*, and in the upregulated group in *La2/+.*

| **Genotype** | **Representation factor of "morphogenetic" genes** |
| --- | --- |
| *clausa* up | **Significantly over-represented Representation factor: 1.5 p < 0.014** |
| *clausa* down | **Significantly under-represented**  **Representation factor: 0.4 p < 0.008** |
| *La2/+* up | **Significantly under-represented**  **Representation factor: 0.6 p < 0.016** |
| *La2/+* down | **Significantly over-represented Representation factor: 1.4 p < 0.011** |
| *pFIL>>miR319* up | **Significantly over-represented**  **Representation factor: 1.3 p < 0.044** |
| *pFIL>>miR319* down | **Not significant**  **Representation factor: 1.2 p < 0.205** |
| *pFIL>>TKN2* up | **Significantly over-represented**  **Representation factor: 2.2 p < 3.954e-28** |
| *pFIL>>TKN2* down | **Significantly under-represented**  **Representation factor: 0.3 p < 1.710e-16** |
| *pFIL>>TKN2*-*SRDX* up | **Not significant**  **Representation factor: 1.1 p < 0.251** |
| *pFIL>>TKN2-SRDX* down | **Significantly over-represented**  **Representation factor: 1.6 p < 9.486e-06** |

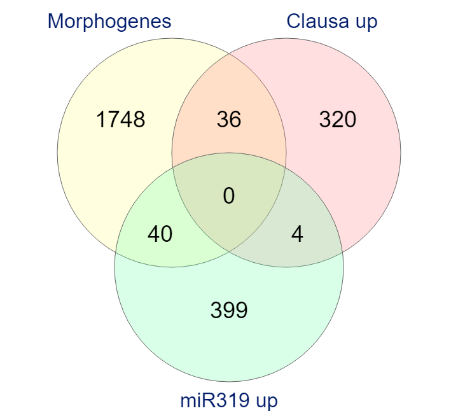
Interestingly, we see that the genes upregulated in *clausa* and *pFIL>>miR319* are different genes, supporting the notion that CLAU and LA operate in parallel pathways.

"Differentiative" genes, as a group, are significantly enriched in the upregulated group in *La2/+*, and in the downregulated group in *pFIL>>TKN2*, *clausa* and *pFIL>>miR319*. They are also significantly depleted in the downregulated group in La2/+ and in the upregulated group in *pFIL>>miR319*, *pFIL>>TKN2*, and, interestingly, *pFIL>>TKN2-SRDX*.

| **Genotype** | **Representation factor of "differentiative" genes** |
| --- | --- |
| *clausa* up | **Not significant**  **Representation factor: 1.4 p < 0.057** |
| *clausa* down | **Significantly over-represented**  **Representation factor: 2.5 p < 1.657e-06** |
| *La2/+* up | **Significantly over-represented**  **Representation factor: 4.0 p < 2.309e-24** |
| *La2/+* down | **Significantly under-represented**  **Representation factor: 0.3 p < 1.772e-04** |
| *pFIL>>miR319* up | **Significantly under-represented**  **Representation factor: 0.7 p < 0.049** |
| *pFIL>>miR319* down | **Significantly over-represented**  **Representation factor: 2.6 p < 2.808e-08** |
| *pFIL>>TKN2* up | **Significantly under-represented**  **Representation factor: 0.4 p < 1.772e-09** |
| *pFIL>>TKN2* down | **Significantly over-represented**  **Representation factor: 2.7 p < 9.404e-33** |
| *pFIL>>TKN2*-*SRDX* up | **Significantly under-represented**  **Representation factor: 0.7 p < 0.027** |
| *pFIL>>TKN2-SRDX* down | **Not significant**  **Representation factor: 1.2 p < 0.150** |

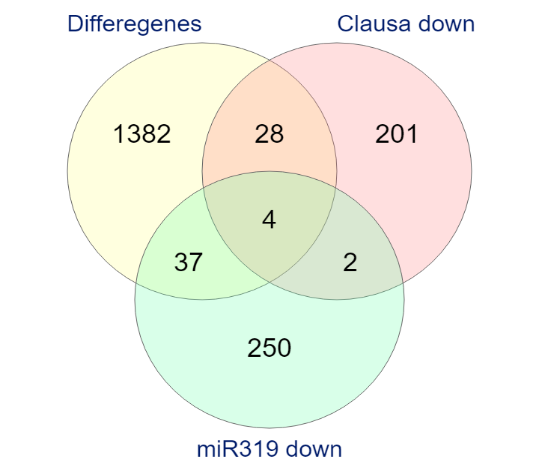
Once again, looking at the specific differentiation genes downregulated in *clausa* and *pFIL>>miR319*, we see very little overlap.
